# Spatiotemporal Microbial Ecoevolutionary Dynamics on the International Space Station

**DOI:** 10.1101/2025.10.28.685245

**Authors:** Megan S. Hill, Anna C. Simpson, Vanessa R. Minnis, Mariana C. Salas Garcia, Alexander Mahnert, Simon Lax, Ella Rushton, Ryan K. Chung, Davis Bone, Ceth W. Parker, Sarah M. Allard, Molly Matty, Kasthuri Venkateswaran, Jack A. Gilbert

## Abstract

This study presents the most comprehensive spatiotemporal analysis of the ISS microbiome to date. Over a seven-year period, 184 surface samples were collected for live/dead metagenomic profiling, phenotypic characterization of antimicrobial resistance (AMR) and virulence among 102 cultured isolates, and metagenome-assembled genome (MAG) analysis to evaluate evolutionary selection pressures. Despite ecological stability, comparative genomics revealed ongoing microevolution through lateral gene transfer and selection for traits (e.g., radiation resistance and biocide tolerance). Importantly, predictions of AMR and virulence frequently misaligned with experimental outcomes, underscoring the need for functional validation. This dataset highlights a stable core microbiome that persists across years, punctuated by localized adaptation and gene flow. The ISS microbiome exemplifies both ecological resilience and microevolutionary innovation that can inform risk management for long-duration spaceflight.

## Introduction

The International Space Station (ISS) is widely considered one of the most extreme human habitats: a hermetically-sealed, micro-gravity outpost that couples chronic radiation, elevated CO₂, and continual human occupation, with rigorous engineering constraints (*1*). Over two decades of occupation have shown that these stressors shape a distinct built-environment microbiome that differs markedly from terrestrial analogues in diversity, metabolism and chemical exposure profiles (*2–8*). ISS surfaces are dominated by human-associated taxa whose composition tracks crew rotation and activity patterns (*9*, *10*), and prolonged spaceflight can remodel both environmental and astronaut-associated microbiota with immunological consequences (*11–14*). Existing surveys using culture-based screens, 16S rRNA gene amplicon sequencing, or spatial analysis using metagenomics and metabolomics have revealed persistent opportunists—including *Staphylococcus aureus*, *Klebsiella pneumoniae,* and *Enterobacter bugandensis*—and catalogued antimicrobial resistance (AMR) determinants and virulence factors enriched in the orbital habitat (*2*, *4*, *6*, *7*, *15*). Subsequent isolate-led pan-genomic and metabolic-modelling efforts underscored the ecological influence of *K. pneumoniae* and other Enterobacteriaceae on bacterial community structure and on inter-kingdom interactions with fungi (*16*). A more recent three-dimensional multi-omic survey of 803 surface samples across nine ISS modules showed that use-pattern-driven, module-specific microbial and chemical signatures place the station at the extreme end of an industrialization gradient, with markedly reduced diversity and enrichment of human-associated taxa compared with terrestrial built environments (*15*). Yet, most investigations have been cross-sectional, limited to short flight series, or have decoupled genotype from experimentally verified phenotype. Consequently, the evolutionary forces governing microbial succession, functional stability and pathogenic potential in confined spacecraft remain poorly resolved, hampering our understanding of ecological dynamics and evidence-based microbial risk management for long-duration exploration missions.

Here, we provide the most extensive spatial and longitudinal analysis of a spacecraft microbiome reported to date. We integrate five years of metagenomic surveys with a seven-year culture collection from the United States Orbital Segment to address three central questions. First, we explore the stability of the viable ISS microbiome using propidium monoazide (PMA) treatment to track viable microbial taxonomic and functional turnover and volatility (*17*), and by quantifying the high-resolution temporal dynamics of antimicrobial resistance (AMR) and virulence-associated genes. Second, we determined whether genomic predictions correspond to realized phenotypes for 102 cultured isolates spanning multiple flights, using *in silico* AMR and virulence prediction with *in vitro* antimicrobial-susceptibility testing and *in vivo* pathogenicity assays. Finally, we determine the signatures of evolutionary selection that emerge in this habitat, by reconstructing and comparing >100 metagenome-assembled genomes (MAGs) with paired isolate genomes. We identify loci under putative positive selection linked to radiation resistance, peroxide resistance, and biocide tolerance, and dissect the evolutionary trajectories of spacecraft-associated bacteria across spaceflight and laboratory contexts. We recently published a genomic and phenotypic characterization of a novel Actinobacterium, *Microbacterium mcarthurae*, isolated from ISS crew quarters during this study (*18*). *M. mcarthurae* showed station-wide enrichment over time, exhibiting resistance to 11 antibiotics (fluoroquinolone-to-β-lactam) and moderate *in vivo* virulence that was not predicted from genotype alone, emphasizing the need for proactive detection of emergent pathogens in spacecraft environments (*18*). This larger study further exemplifies this need.

Our integrated eco-evolutionary framework reveals that, despite pronounced short-term fluctuations in microbial DNA signatures driven by crew exchange and module use, the ISS community exhibits a resilient core of viable bacteria whose functional repertoire remains remarkably conserved, highlighting stable selection for specific species. Comparative genomics and metagenomic reconstructions further showed that, within this stability, the ISS microbiome undergoes active diversification with loci under positive selection linked to desiccation tolerance, radiation resistance, and host interaction, as well as lateral gene transfer events that disseminated stress-response functions across taxa. At the same time, by integrating genomic and culture-based assays, we found that in silico predictions of antimicrobial resistance and virulence factors often failed to capture actual phenotypes, with resistant genotypes lacking expression and, conversely, ostensibly benign strains exhibiting pathogenic traits in vitro. Together, these results demonstrate how the ISS microbiome simultaneously sustains ecological stability and microevolutionary variation, while underscoring the need to validate sequence-based predictions with experimental assays for accurate risk assessment.

By coupling time-series metagenomics with experimental phenotyping, our study demonstrates how confined built environments foster both ecological stability and rapid microevolutionary change, and it provides an empirical basis for predictive models of microbial risk in future lunar and Martian habitats. These findings advance our understanding of microbial selection beyond Earth, highlight the need for culture-independent surveillance thresholds tailored to spaceflight conditions, and furnish a benchmark dataset for designing targeted mitigation strategies that safeguard crew health on forthcoming deep-space missions.

## Results

Samples were collected from the International Space Station (ISS) over a seven-year period across three missions (2015–2022) from eight surfaces distributed across five modules (Fig. 1), yielding a total of 184 experimental samples. For Mission 3, five consecutive days of sampling were performed for each of the three flights (Flight 9–11), with aliquots processed for qPCR, metagenomics ± propidium monoazide (PMA), and microbial culturing. From cultured isolates, expressed phenotype was compared to genotypic prediction for pathogenic traits, using *in silico* testing of 102 taxonomically diverse isolates for antimicrobial resistance (AMR) (n = 67) and virulence (n = 79) assays (n = 44 strains shared across experiments).

**Fig. 1.**
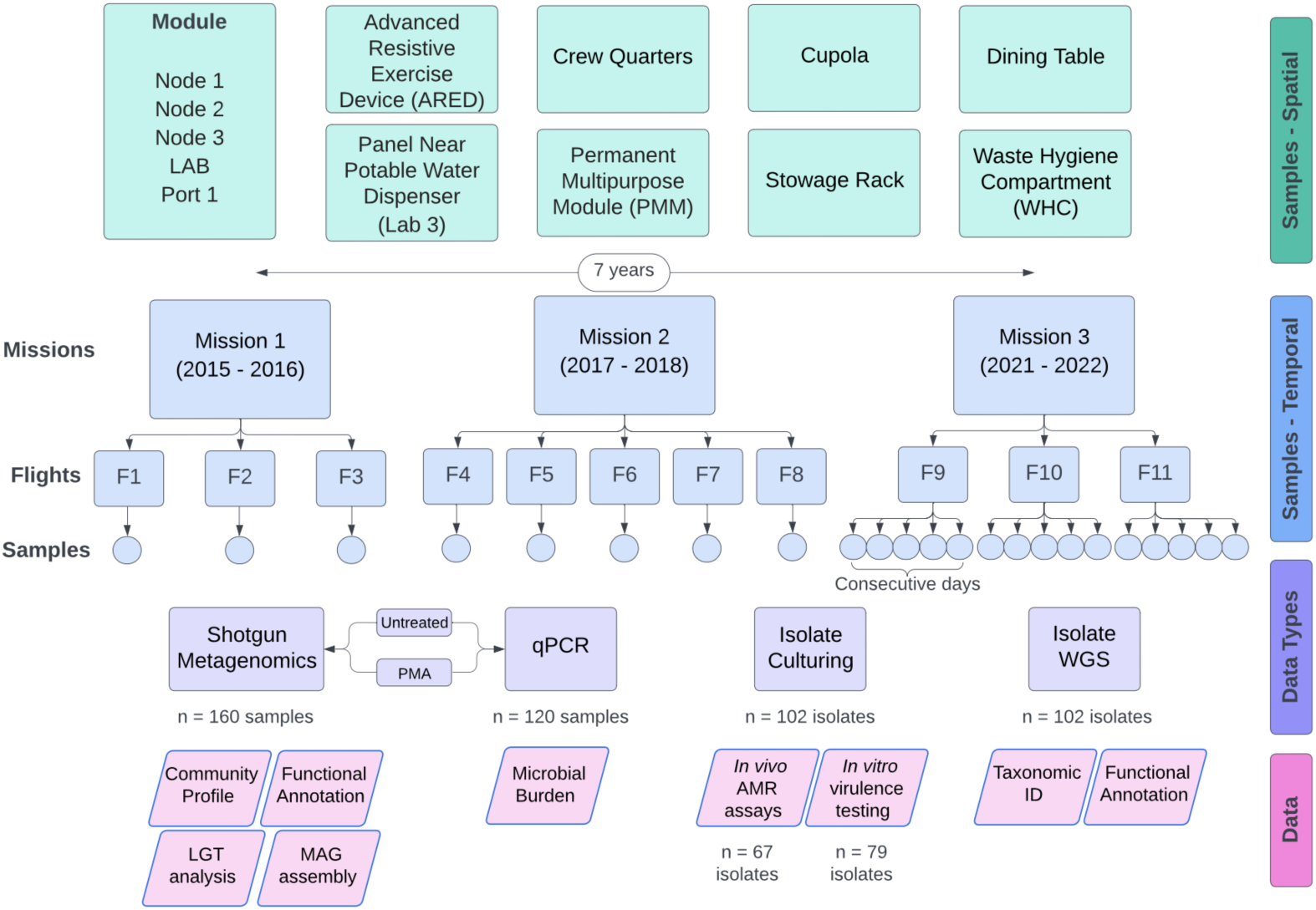
Project overview. Hierarchy of spatiotemporal sample collection, including eight surfaces from five ISS modules, as well as sample types and data generated. Due to differences in storage temperature requirements or delay in sample return, samples from Mission 1 and Flight 8 of Mission 2 were excluded from community-based metagenomic analyses.

### Microbial abundance and community structure

qPCR analyses revealed that bacterial gene copies (av. ∼220,000 16S rRNA copies/sample) significantly exceeded fungal gene copies (av. ∼130,000 ITS copies/sample) in untreated (not PMA-treated) samples (p<0.001, log₁₀ ng/sample, Wilcoxon). However, when restricted to viable communities (PMA-treated), fungi and bacteria had similar gene copy numbers, albeit with significantly greater ITS copies (16S av. ∼30,000; ITS av ∼32,000; p<0.001, log₁₀ ng/sample, Wilcoxon), driven by high ITS copy numbers recovered from a subset of permanent multipurpose module (PMM) and stowage rack samples (Fig. S1). Notably, these locations represent areas of similar usage and material. Overall, microbial gene copy numbers were consistently greater on the PMM and stowage rack than on other surfaces (Fig. S1), though viable bacterial abundance did not differ significantly among surfaces (Table S1).

Bacterial community composition predominantly derived from human occupants and was shaped by both viability and surface type (fungal community structure was not assessed in this study). Untreated and PMA-treated bacterial beta diversity differed modestly between missions (p=0.003, Aitchison distance, PERMANOVA, R²=0.01; Fig. 2a). Surface explained more variation in beta diversity than did the mission or flight (untreated: R²=0.18, p<0.001; PMA-treated: R²=0.09, p<0.001; Fig. 2b). The waste and hygiene compartment (WHC) bacterial community was the most distinct among untreated samples (p<0.001 for all surfaces) and was enriched in 38 taxa that are primarily of fecal origin, including *Fusicatenibacter saccharivorans*, *Blautia massiliensis*, *Faecalibacterium prausnitzii*, *Mediterraneibacter faecis*, and *Eubacterium rectale* (>3 log-fold increases; Fig. 2c). However, these differences disappeared in PMA-treated sample analysis, suggesting poor viability of gut-associated taxa on ISS surfaces (Fig. 2c). In contrast, the viable community was most distinct on the PMM (p<0.05 for all comparisons, Aitchison Distance, PERMANOVA; Fig. 2d).

**Fig. 2.**
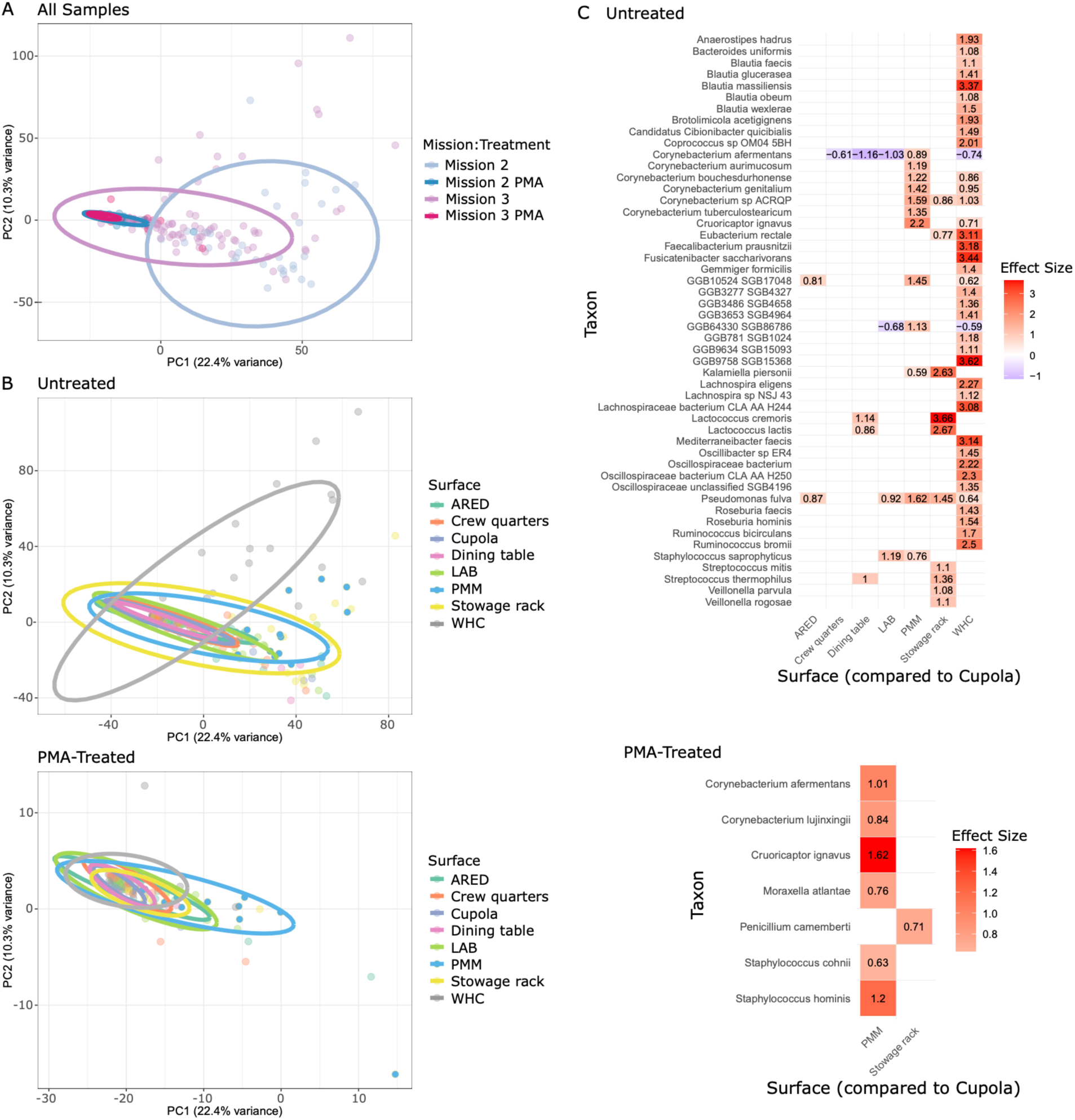
Community-level differences among ISS sample types. Ordinations are based on Aitchison distance from CLR-transformed SBG tables. (**A**) Temporal variation across all experimental sample types collected during Missions 2 (archival) and 3 (new), grouped by PMA treatment. (**B**) Spatial variation among Mission 3 samples by surface, with the WHC showing the highest variability in untreated communities and the PMM showing the highest variability in viable (PMA-treated) communities. (**C**) Differential abundance analysis of surfaces relative to the Cupola port panel (ALDEx2) mirror beta-diversity analysis, where the WHC shows the strongest enrichment of fecal-associated taxa in untreated samples, an effect absent in PMA-treated samples.

### Temporal and Spatial Dynamics

Community volatility was consistently greater in untreated than in PMA-treated samples, with WHC showing the highest variability, as a result of non-viable gut-associated bacteria (mean within-group Aitchison distance (± SEM), Fig. S2a). Volatility decreased at shorter intervals; i.e., samples collected within missions (months apart) were more stable than those collected between missions (years apart) (Fig. S2a,b). Exceptions were the PMM and stowage rack, where viable communities remained stable across all time periods (Fig. S2a).

From PMA-treated Mission 3 metagenomes, 251 unique species were identified, with *Cutibacterium acnes* (36%), *Ralstonia pickettii* (17%), *Microbacterium enclense* (13%), and *Massilia timonae* (4%) dominating (Fig. S3a). Differential abundance analysis identified 15 taxa with significant changes in relative abundance between missions, with *R. pickettii*, *M. enclense*, and *M. timonae* showing the greatest magnitude of change (p≤0.001, effect sizes 0.73–1.27). Random Forest models confirmed their predictive power for distinguishing Mission 2 vs. Mission 3 (Fig. S3b). Interestingly, these species were only enriched in specific flights; e.g., *R. pickettii* in Flight 11, and *M. enclense* and *M. timonae* in Flights 9 and 10, suggesting that human occupancy or activity could be driving small differences over time (Fig. S3b).

Skin-associated taxa remained consistently abundant. On average, 40% of bacterial reads mapped to 25 common skin bacteria (down from 59.1% in Mission 2; p=0.013, Wilcoxon; Fig. S3c). Surfaces with the greatest enrichment of skin bacteria included the panel used for materials science research (LAB) (59%), the advanced resistive exercise device (ARED) footpad (46%), and Cupola port wall (43%), with the stowage rack having the lowest average proportion (23%) (Fig. S3d). Despite differences in usage, proportions did not differ significantly among surfaces (p>0.05, Kruskal–Wallis). This suggests that the viable ISS bacterial communities are ecologically stable over short timescales, with subtle changes due to species introductions between missions.

### Functional Potential of the Viable Microbiome

Metagenomic data from 160 PMA-treated samples revealed no significant differences in AMR gene proportion between missions (p=0.138, Wilcoxon). Flight-level variation was modest, with significantly decreased AMR prevalence over time (p=0.01–0.015; Fig. 3a). No differences were observed between consecutive sampling days (Table S2). Spatially, the PMM showed AMR enrichment relative to the ARED (p=0.013, Wilcoxon), crew quarters (p=0.006), Cupola (p=0.001), dining table (p<0.001), and stowage rack (p<0.001) (Fig. 3a), and the PMM was the only surface that had a significant increase in the proportion of ARGs between missions (p=0.044). Resistance profiles (21 pathways) were stable across missions and surfaces (p>0.05, Wilcoxon with BH correction, Fig. S4) and dominated by MLS genes (57,818 CPM), with sulfonamide genes as the least abundant (1 CPM). Virulence factor (VF) proportions also remained stable across missions (p=0.25, Wilcoxon; Fig. 3b), flights, and consecutive sampling days (Table S2). No VFs were observed from Cupola, stowage rack, or WHC. The LAB and PMM carried the highest average VF proportions (0.062–0.072), while crew quarters had the lowest (0.001–0.002). The PMM’s physical location changed between Missions 2 and 3 (Port 1 → Deck 1), which coincided with altered crew usage (introduction of drying clothes, sponge baths, foot-bar use), potentially influencing ARG and VF prevalence.

**Fig. 3.**
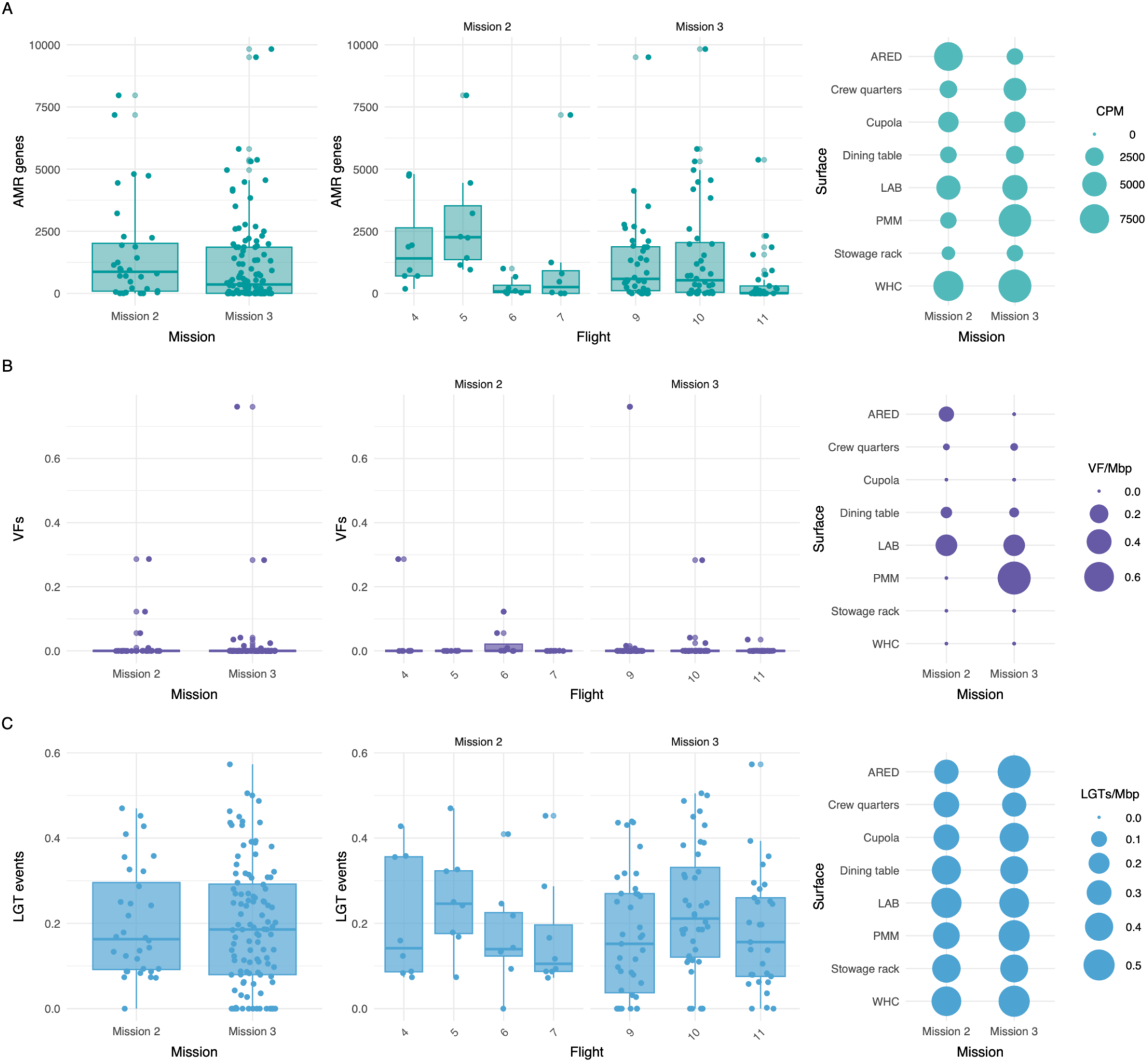
Proportion of antimicrobial resistance (AMR) genes, virulence factors (VFs), and lateral gene transfer (LGT) events at spatiotemporal scales on the ISS. Comparisons include differences between missions, flights within missions, and individual surfaces over time, based on Wilcoxon rank-sum tests, using a Benjamini-Hochberg correction. Results suggest spatiotemporal stability of genes associated with pathogenicity, as well as transfer rates of genetic material among bacteria (**A**) ARG abundance was normalized by millions of sequences per sample (CPM). No statistically significant differences were observed between missions, overall or for individual surfaces over time. Modest differences observed between flights. (**B**) VFs were normalized by contig length (VFs/Mbp). No statistically significant differences were observed across all comparisons. (**C**) LGT events were normalized by LGT events/Mbp. No statistically significant differences were observed across all comparisons.

Lateral Gene Transfer (LGT) events, quantified from untreated samples (to preserve potential sharing of non-cellular DNA), revealed 1155 events between 517 taxa (Mission 2) and 1670 events between 643 taxa (Mission 3). *Campylobacter jejuni* (4.63%; Mission 2) and *Staphylococcus epidermidis* (3.47%; Mission 3) had the greatest proportion of LGT events. Transfers were biased toward related taxa, especially among staphylococci. LGT prevalence did not differ between missions (p=0.24, Wilcoxon) or flights (Table S2), but was surface-specific, with the WHC showed the greatest rates (0.29 LGTs/Mbp), exceeding the crew quarters (p=0.005, 0.06 LGTs/Mbp) and dining table (p=0.016, 0.111 LGTs/Mbp; Fig. 3c). Therefore, while functional traits (AMR, VF) are stable over time, they did show localized enrichment due to crew activities, and frequent LGT events suggest evolutionary plasticity within a resilient resistome and virulome.

### Linking Genotypes to Phenotypes

Across three missions, 67 bacterial isolates were tested against 18 antibiotics, representing four antibiotic classes (Table S3). Genomic predictions frequently misaligned with phenotypes, with 76% of strains showing discrepancies (Fig. 4a). Thirty strains carried resistance genes but were not resistant (false positives), while 31 showed phenotypic resistance without corresponding genes (false negatives; Table S3). False negatives spanned multiple genera and surfaces, with *Enterococcus faecalis* and *Staphylococcus saprophyticus* exhibiting broad β-lactam resistance (Fig. 4a). Only azithromycin and levofloxacin resistance were perfectly predicted genomically.

**Fig. 4.**
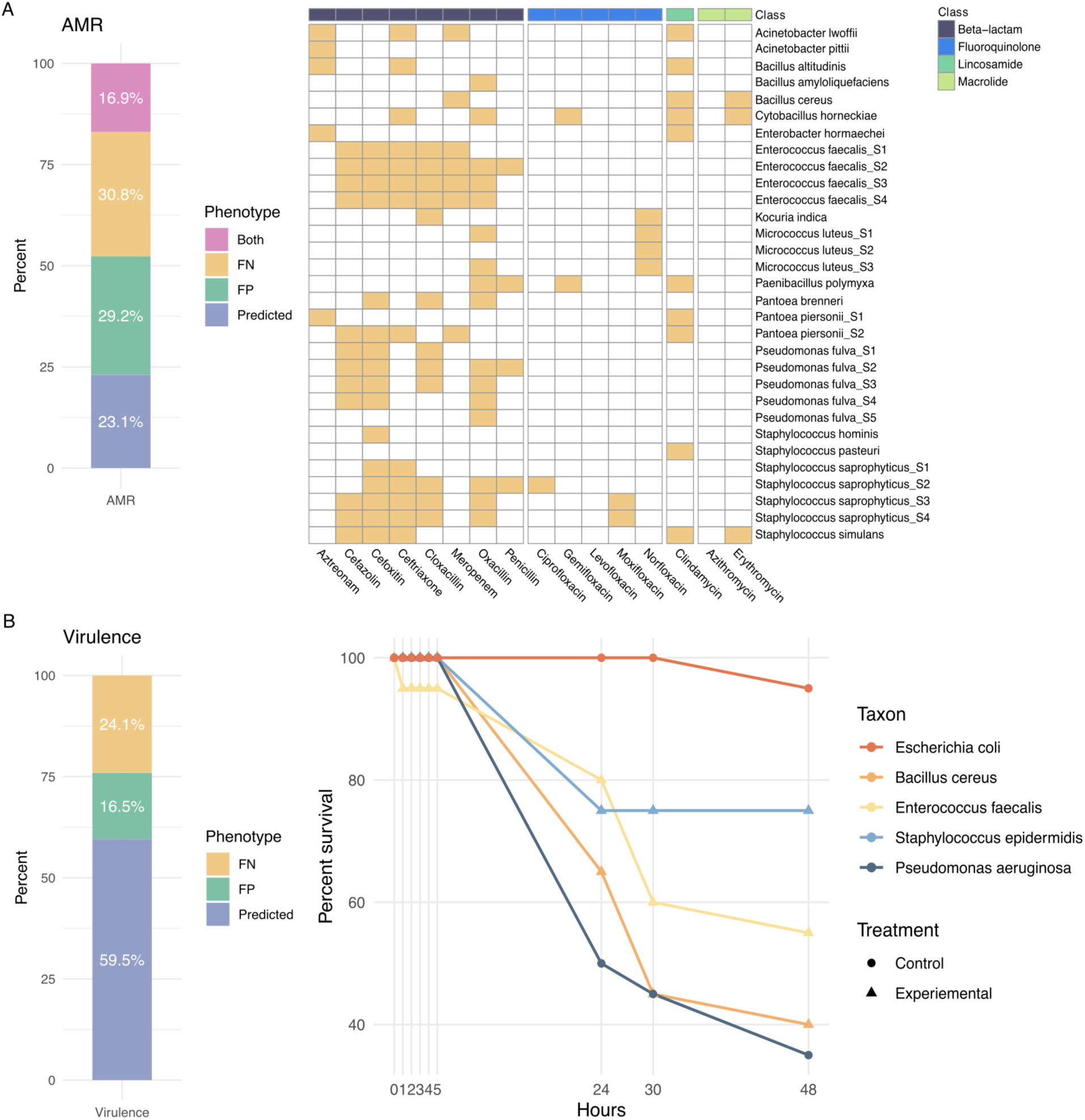
Antimicrobial resistance (AMR) and virulence phenotypic results. (**A**) AMR results. Left panel represents the percent of strains that had a false negative (FN), false positive (FP), or predicted phenotype, based on genomic data. Strains that exhibited both FN and FP results (different antibiotics) are classified as ‘both’. Right panel represents strains that exhibited a false negative phenotype (i.e., resistance was present but not predicted) across all missions. All strains were tested using Kirby-Bauer assays against 18 antibiotics that spanned four antibiotic classes (n = 67 strains, XX species). Multiple strains of the same species denoted with suffixes (e.g., *S1, S2*). (**B**) Virulence results. Left panel represents the percent of strains that had a FN, FP, or predicted phenotype, based on genomic data. Right panel represents strains classified as highly virulent across all missions. Virulence was tested in a *C. elegans* model (n = 79 isolates; 22 species), where percent survival was averaged across replicates, prior to calculation of differences in mean lifespan and survival over time, compared to *E. coli* OP50 (EC; negative control) and *P. aeruginosa* PA14 (PA; positive control). Strains were considered highly virulent if the rate of killing was equal to or greater than that of *P. aeruginosa* (no statistical difference from *P. aeruginosa* or significantly higher than *P. aeruginosa*).

Virulence assays in *C. elegans* (79 isolates) revealed 47 avirulent, 29 intermediate, and 3 highly virulent strains (Table S4, Fig. 4b). High-killing strains included *Bacillus cereus* (Mission 1), *S. saprophyticus* (Mission 1), and *Pseudomonas granadensis* (Mission 2) (Fig. 4b). Genomic VF content poorly predicted outcomes, with 13 false positives (>20 VFs but avirulent/intermediate phenotype) and 19 false negatives (no VFs but intermediate/highly virulent phenotype). *Acinetobacter pittii* strains had a high rate of false negatives (n=4/5), followed by *Staphylococcus* spp. (7/10). Therefore, genomic predictions often misrepresented realized phenotypes and phenotypic assays reveal potential hidden risks (e.g., β-lactam resistance, unexpected virulence), underscoring the need for functional validation.

### Eco-Evolutionary Signals

From Mission 3, 36 high-quality MAGs (95±5% completeness, 2±2% contamination, from 252 dereplicated genome bins) were assembled to facilitate strain-level insights (Fig. 5a). The covered genome fraction between MAGs and assemblies averaged 3.9±17.2% from 235 samples, reaching a maximum of 100% for the WHC. *C. acnes* MAGs covered up to 47.36% of all reads with the greatest mapping at the ARED during flight 9 (68.72%). *C. acnes* (41.6±43.4% proportion per sample assembly) and staphylococci were the most prevalent (>10% proportion per sample assembly). Predicted growth rates averaged 0.08±0.3, differing by location (q<0.001, FDR-corrected Kruskal-Wallis rank sum tests), with *C. acnes* showing potential for growth on 75% of surfaces. Average growth optima suggested adaptation to ISS conditions (pH: 7±1, NaCl: 4±3%, temperature: 31±3°C), with significant oxygen tolerance differences across families (q<0.001, FDR-corrected Wilcoxon rank sum test and Fisher’s exact test). AMR and VF profiles yielded 129 hits. *S. epidermidis* (WHC) had the highest antimicrobial potential, carrying 22% of AMR genes, while two *Neisseria* species (stowage rack) carried 90% of VF hits. Individual strain profiling revealed significant SNP-level differences between flights (p=5.8×10⁻⁸–1.3×10⁻²¹, pairwise Wilcoxon rank sum tests), surfaces (ARED, Lab, WHC: p = 1.0–4.1×10⁻²), and specific taxa (Actinomyces sp900323545 vs. *Agathobacter rectalis*: p=3.4×10⁻⁴).

**Fig. 5.**
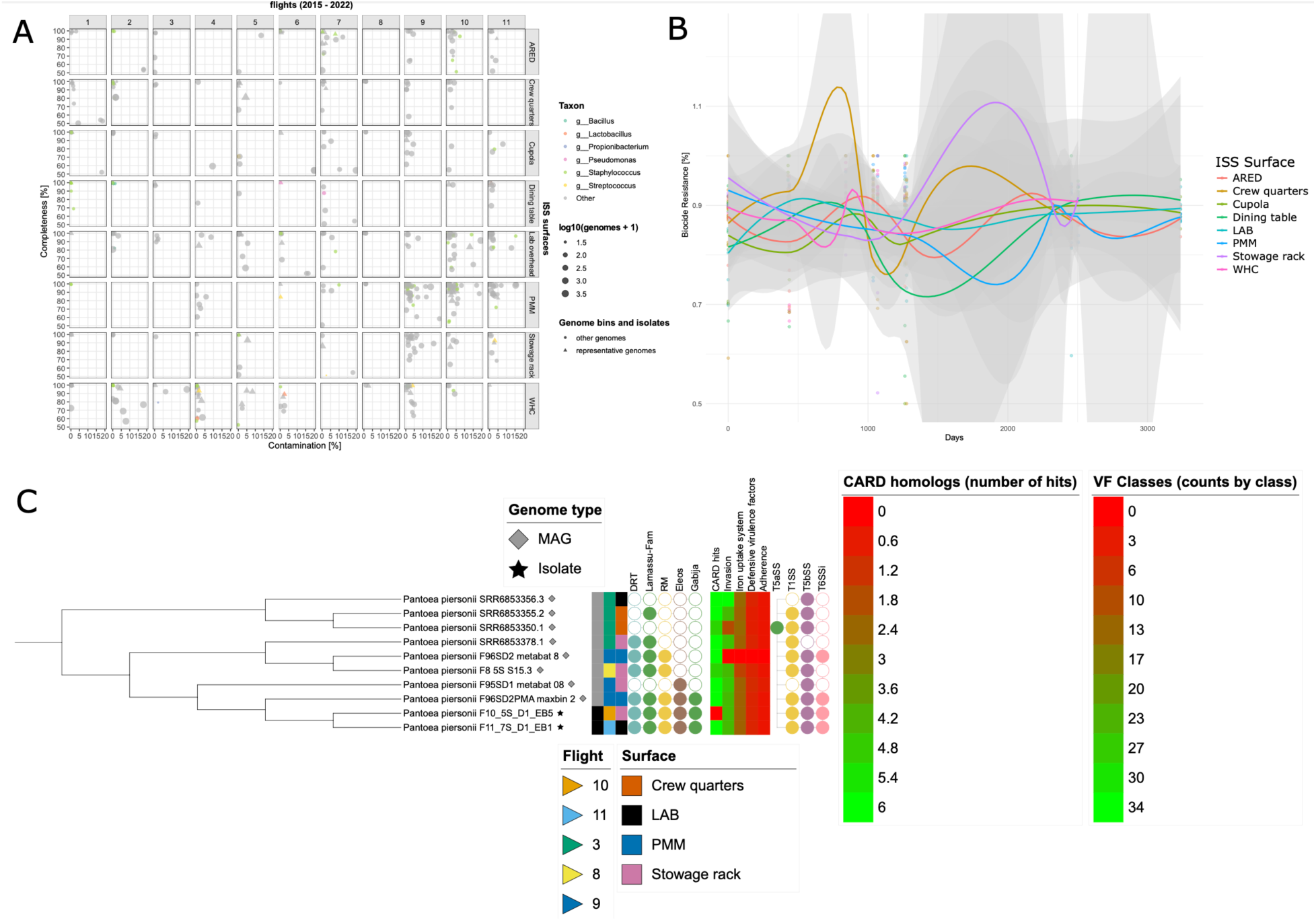
Metagenome-assembled genome (MAG) results. (**A**) Bubble plot showing the spatial (ISS surfaces) and temporal (flights) distribution of all genome bins and isolates (>50% completeness, <20% contamination) from Missions 1, 2, and 3. Representative genomes are displayed as triangles, all others as circles. Sizes are scaled according to the log10 of the number of retrieved genomes. Most prevalent taxa in the dataset are colored on genus level. (**B**) Line plot based on volatility analysis showing relative abundance of biocide resistance of genome bins and isolates on ISS surfaces along the entire sampling timeframe according to DRAM2 formatted databases. (**C**) Interactive midpoint rooted rectangular tree of selected annotated *Pantoea piersonii* genomes visualized with iTOL. Annotations from left to right: genome type, flight, ISS surface, defense systems, CARD homologs, virulence factor classes, and macromolecular systems.

Comparing MAGs from Mission 3 to genome bins from all three missions revealed that certain highly abundant species, including *C. acnes*, *Staphylococcus hominis*, *S. epidermidis*, and *Pantoea piersonii*, displayed greater genome coverage (Fig. 5a). From a dereplicated set of 99 high-quality MAGs, metabolic annotation and feature volatility analysis, focused on carbon utilization and engineered systems, highlighted glycosyl transferases and biocide resistance as the most prominent functional traits. While temporal or surface-specific trends were generally variable, the PMM consistently harbored greater relative proportions of biocide resistance compared to other surfaces (Fig. 5b).

Additionally, we created a representative spatiotemporal genome catalog of the ISS environment, based on 99 high-quality dereplicated MAGs from 673 genome bins that were dominated by *Staphylococcus* spp., resulting in an expanded database of 114 representatives. Most genomes belonged to *C. acnes* (73), *P. piersonii* (14), *S. hominis* (14), and species within the *Corynebacterium* genus (13). This resulted in a slight shift in the most abundant species, including *C. acnes* (73), *Pseudomonas fulva* (20), diverse *Staphylococcus* species (n = 18 *S. hominis*, 16 *S. saprophyticus*, and 15 *S. epidermidis*), *P. piersonii* (16), *Acinetobacter pittii* (13), and the *Corynebacterium* genus (13). Based on *in vitro* and *in vivo* experiments and the coverage across all sampling events, we selected two isolates and eight MAGs of *P. piersonii* (growth rate prevalence on all sampled surfaces: 7.2%; pH optimum 6.8; salinity optimum 2.2%; temperature optimum 31.5 °C) as a representative example for an in-depth comparative analysis including gene synteny.

The case study of *P. piersonii* genomes from all missions revealed two clusters (cultured vs. MAGs, Fig. 5c). Since genomes did not cluster according to sampling event, we expected a stable genome content over the entire sampling timeframe (0 – 2495 days). Of these, we calculated gene phyloprofiles for two isolates (F10_5S_D1_EB5, F11_7S_D1_EB1) compared to other ISS-associated genomes (excluding genes common to all other *Pantoea*). F10_5S_D1_EB5 showed lower gene conservation and fewer shared homologs, indicating greater genomic divergence, which might reflect higher rates of adaptation through gene gain or loss (e.g., a potential antibiotic biosynthesis monooxygenase and a copper resistance protein). Genome plasticity analysis revealed that F10_5S_D1_EB5 contained more putative horizontally acquired genomic islands and unique regions than F11_7S_D1_EB1, consistent with its greater genomic divergence. Both isolates harbored a high-scoring candidate island near a tRNA gene, suggesting integration of mobile elements.

Comparative synteny analysis showed that while ISS-associated *P. piersonii* genomes generally maintained conserved gene order compared to broader *Pantoea* species, synteny varied between isolates, even when sampled close in time and space, highlighting micro-diversity in genome organization. Analysis of gene fusion and fission events revealed frequent structural changes (>30% fissions, >10% fusions), though most lacked functional relevance. Only a few transport- and resistance-related events suggested possible adaptive traits to the ISS environment. Some examples include hits for multidrug, radiation, fosfomycin, copper, and peroxide resistance that could potentially be related to the specific ISS environment, though these achieved only low scores. Additionally, we observed *P. piersonii* genomes share a relatively conserved core of ∼3,000 gene families, compared to an open pangenome analysis that exceeded 30,000 families, underscoring the narrower but more cohesive gene repertoire of the ISS isolates. CRISPR, prophages, and defense elements were abundant (average 5±2 hits per genome). AMR and virulence signatures were consistent across genomes; most of which belong to the class of an antibiotic efflux mechanism (87%), with an average of six AMR genes detected per genome. Potential virulence genes were in a similar range (average: 75±7 hits per genome), primarily associated with flagellar motility and siderophore production (e.g., enterobactin, aerobactin).

Despite ecological stability, the ISS microbiome undergoes microevolution, acquiring adaptive traits (e.g., radiation resistance, peroxide resistance, biocide tolerance) through LGT and selection.

## Discussion

This study provides the most comprehensive spatiotemporal analysis of the ISS microbiome to date. We collected 184 surface samples across three missions (over 7 years) and sequenced them with and without viability selection (PMA treatment), while also obtaining 102 bacterial isolates for genome sequencing and phenotypic assays. This integrated dataset far surpasses previous ISS surveys in temporal resolution and breadth, offering an unprecedented resource for understanding microbial ecology associated with spacecraft. Earlier culture-independent studies had hinted at the persistence of certain taxa and AMR genes on the station and noted differences between total vs. viable microbial communities (*5*, *6*, *19–21*). Building on that foundation, our work couples high-frequency longitudinal sampling with sequencing and culturing techniques, enabling a data-driven view of how these microbial communities establish, endure, and evolve under spaceflight conditions. We present paired pathogenic profiling of ARGs and VFs, analyses of LGT events from metagenomic data, experimental verification of phenotypic expression of pathogenic traits for the largest set of cultures isolates tested to date, and direct comparisons of cultured isolate genomes against MAG-based reconstructions.

Consistent with studies from early space habitats, such as Skylab-4 and the Mir space station, we found abundant Gram-positive bacteria, particularly *Staphylococcus* and *Micrococcus* species (*22*, *23*). Further, on the ISS, traditional culture-based techniques and culture-independent metagenomic analyses have consistently recorded a microbial composition predominantly associated with humans and showed a noticeable loss of diversity compared to environments on Earth (*5*, *15*, *20*). Despite the station’s extreme environmental pressures and routine disturbances, the viable (PMA-treated) community demonstrated notable ecological stability. On short timescales (day-to-day within each mission), community composition remained largely unchanged, suggesting a resilient core microbiota under strong stabilizing selection. Transient perturbations (e.g., crew turnover or module-specific activities) caused only modest, temporary shifts in DNA profiles, and these did not fundamentally alter the active community. This stability indicates that, once established, the ISS microbial ecosystem is buffered against minor fluctuations, maintaining a conserved functional repertoire over weeks to months. Such findings align with the notion of *in situ* selection for well-adapted species in the space station environment. At the same time, clear spatial differentiation was observed in certain areas with microbial profiles varying by location, driven by surface usage patterns and local environmental conditions. For example, the toilet (WHC) vs. storage and personal hygiene locations (PMM) showed usage-specific microbial compositions, reminiscent of those seen in Earth-based built environments (*24*, *25*).

Over longer periods, we detected gradual temporal changes, indicating that the ISS microbiome is not static. Notably, some taxa exhibited significant shifts in abundance across the seven-year span. For example, an actinobacterial lineage that we isolated (recently described as *M. mcarthurae*) showed a station-wide increase in proportion over this time period, accompanied by a set of previously undescribed AMR and virulence genes (*18*). Such successional trends suggest that prolonged habitation and repeated human influx continue to shape community structure, allowing certain opportunistic or stress-tolerant organisms to stably adapt. Importantly, however, even these long-term changes unfolded within the framework of a stable core community – no unchecked pathogen blooms were observed and the core resistome and virulome remained stable over time and space, despite frequent exchange of genetic material through LGT events. This balance of stability and slow succession underscores that the ISS microbiome, while constrained and predominantly human-sourced, is still an adaptive ecosystem. The station’s isolation on the extreme end of the built-environment spectrum thus leads to a depauperate but dynamically maintained community, one that can inform how microbial communities might behave in future closed habitats.

By integrating genomic and culture-based analyses, we were able to directly test how well genotypes predict phenotypes in this environment. We found that *in silico* predictions of AMR and virulence was a poor predictor of phenotype. Approximately three-fourths of isolates (n=67) that were carrying resistance genes did not exhibit the expected drug resistance *in vitro*, and one-fourth of strains (n=79) without known virulence factors showed the capacity to cause disease in our model assays, including known opportunistic pathogens (e.g., *Acinetobacter pittii*). These false positives and negatives highlight the inherent complexity of inferring function from genetic content alone. For example, although no genes were identified that would confer the observed resistance, *E. faecalis* is known to have intrinsic resistance to β-lactam antibiotics due to the presence of low-affinity penicillin-binding proteins, and *S. saprophyticus* often produce β-lactamase enzymes, which can render antibiotics within this class ineffective (*26*, *27*). Additionally, one of the highest killing strains (*P. granadensis*) is generally considered as safe in Earth-bound environments, as it is a soil-associated bacterium; however, there have yet to be any studies conducted in the context of human health (*28*). These results, while supporting inferences made from earlier ISS studies that relied only on sequencing data, suggest that such reliance may overestimate risk if dormant genes are counted, or miss emergent traits not captured in databases. Therefore, phenotyping analysis *in vitro* and *in vivo* allowed us to flag discrepancies and discover novel traits. Overall, this represents the largest genotype–phenotype correlation effort yet undertaken in a spaceflight context, bridging a gap left by prior studies. It demonstrates the value of validating genomic insights with cultivation and bioassays to achieve a more accurate risk assessment and understanding of microbial behavior in spacecraft.

Comparative genomics and metagenomic reconstructions further revealed that the ISS microbial community, despite its stability, is undergoing microevolution. We recovered 67 representative MAGs and compared them with their cultured counterparts, finding evidence of genetic diversification and LGT among surface-associated bacteria. Closely related strains of *P. piersonii* isolated from different surfaces shared mobile genetic elements, including plasmids and gene cassettes encoding stress-response functions. This indicates active gene flow across taxa in the ISS microbiome. These LGT events suggest that even in an isolated ecosystem, bacteria can innovate by swapping genes, potentially bolstering their survival under space-specific stresses. In parallel, we identified several loci under putative positive selection linked to adaptative traits, such as desiccation tolerance, biofilm formation, radiation resistance, and host interaction. These genetic signatures of selection point to the pressures that are somewhat unique to the ISS (e.g., low water availability, high radiation, and frequent cleaning regimes) that are shaping microbial evolution. Taken together, the LGT events and adaptive mutations portray an ecosystem exhibiting “rapid microevolutionary change” alongside its ecological stability.

In summary, our findings show that the microbial communities on the ISS achieve a balance between persistence and adaptation. The viable bacterial population is anchored by a stable core that preserves key functions across days and even years, even as subtle shifts and genetic innovations accumulate. This duality of ecological steadiness amid ongoing evolution provides insights into how microbes adapt to the rigors of long-term space habitation. Notably, we did not observe any specific virulence outbreaks or rampant pathogen proliferation; instead, the dynamics resemble those in terrestrial built environments, albeit in an exaggerated form (extremely low diversity and primarily human origin). The presence of opportunistic pathogens (e.g., *S. aureus*, *K. pneumoniae*) on ISS surfaces remains a concern, but our data suggest these organisms persist as part of a structured community, rather than expanding uncontrollably. This nuanced understanding can refocus efforts toward managing the space station microbiome as an ecosystem, using ecological principles rather than only hazard-based metrics.

The technical and conceptual advances of this work offer valuable directions for future research and habitat design. By capturing high-resolution temporal changes and linking genes to functions, we have established a baseline for how a closed-system microbiome behaves over time. This dataset will be a benchmark for testing predictive models of microbial dynamics in space and for developing next generation monitoring tools. For instance, our results support the notion that routine metagenomic sequencing (with viability markers) could complement or enhance traditional culture-based surveillance on spacecraft. Early-warning thresholds might be devised based on functional shifts in the core community rather than total microbial counts alone. Additionally, understanding the stable core’s composition and requirements could inform probiotic or engineering strategies to cultivate a beneficial microbiome in future stations, as opposed to over-sanitization. Therefore, this longitudinal study moves us toward evidence-based microbial management for long-duration spaceflight, and underlines that maintaining astronaut health is not just about eliminating pathogens but about guiding the microbial ecosystem toward a balanced, resilient state. By illuminating how microbes spatially and temporally organize on the ISS, we have laid the groundwork for designing mitigation strategies and habitats that harness microbial stability while curbing undesired evolutionary trajectories. This knowledge will be crucial as humanity prepares for the next era of exploration beyond Earth.

## Acknowledgements

We would like to thank our payload team, including Gwo-Shing Sun, Liz Pane (Payload Manager), and America Reyes Wang (PD Ops Lead). We acknowledge the astronauts that collected Mission 3 samples; Raja Chari, Akihiko Hoshide, Shane Kimbrough, Matthias Maurer, and Megan McArthur, as well as the implementation team at NASA Ames Research Center. Additionally, we thank Olga Zaborina and Alex Zaborin at the University of Chicago for their additional advice in the development of virulence testing protocols, as well as for providing a culture of *P. aeruginosa* PA14. This publication includes data generated at the UC San Diego IGM Genomics Center utilizing an Illumina NovaSeq 6000 and a NovaSeq X Plus that were purchased with funding from a National Institutes of Health SIG grant (#S10 OD026929).

## Funding

Space Biology Grant 80NSSC19K1604.

## Author Contributions

Conceptualization: KV, JAG; Methodology: MSH, ACS, VRM, MCSG, AM, SL, ER, RKC, DB, MM, KV, JAG; Investigation: MSH, ACS, VRM, MCSG, AM, SL, RKC, DB, CWP; Visualization: MSH, ACS, AM, SL; Funding acquisition: KV, JAG; Project administration: MSH, ACS, MCSG, SMA, KV, JAG; Supervision: MSH, ACS, MCSG; Writing – original draft: MSH, ACS, VRM, AM, SL, ER, JAG; Writing – review & editing: MSH, ACS, VRM, MCSG, AM, SL, ER, RKC, DB, CWP, SMA, MM, KV, JAG.

## Competing Interests

The authors declare no competing interests.

## Data and Materials Availability

Data generated during this study and associated metadata are publicly available in NASA’s Open Science Data Repository (OSDR), including isolate genomes from Mission 1: https://osdr.nasa.gov/bio/submission/study/OSD-730, isolate genomes from Mission 2: https://osdr.nasa.gov/bio/submission/study/OSD-731, isolate genomes from Mission 3: https://osdr.nasa.gov/bio/submission/study/OSD-732, and assembled Mission 3 metagenomes: https://osdr.nasa.gov/bio/submission/study/OSD-733. Raw metagenomic data are also available in the NCBI SRA database (Project #: PRJNA1328788), and preprocessing and analysis code are available on GitHub at: https://github.com/hillms/MicrobialTracking-3

## Supplementary Materials

Materials and Methods

Figs. S1 to S4

Tables S1 to S6

References (1-65)

## Notes

### Competing Interest Statement

The authors have declared no competing interest.

