## Supplemental Materials for "Spatiotemporal Microbial Ecoevolutionary Dynamics on the International Space Station"

**The PDF file includes:**

**Materials and Methods**
**Figs. S1 to S4**
**Tables S1 to S6**
**References (*1–65*)**

### Materials and Methods

Sample collection

All samples were collected from the ISS over a seven-year period, and include previously collected samples from the Microbial Tracking-1 (Mission 1) (2015 - 2016, n = 24) and MT-2 missions (2017 - 2018, n = 40) (henceforth referred to as Missions 1 and 2), as well as newly collected Mission 3 data (2021 - 2022, n = 120). More details about the collection methods for Missions 1 and 2 have been previously described and can be found in (*1*, *2*), but briefly, samples were collected once per flight (Mission 1: n = 3 flights, Mission 2: n = 5 flights) from eight unique surfaces, including the foot platform of the Advanced Resistive Exercise Device (ARED), the Crew Quarters-2 Bump-out exterior aft wall, the port panel of the Cupola, the dining table, panel of the Materials Science Research Rack 1 (LAB), the Permanent Multipurpose Module (PMM), the zero G stowage rack, and the forward side panel wall of the Waste and Hygiene Compartment (WHC). Surfaces were distributed across four modules; Node 1, Node 2, Node 3, and the U.S. Lab (Table S5).

Newly collected samples were collected from the same eight surfaces, with the exception of the PMM. Prior to Mission 3, the PMM sample site was moved to a different location within the same module, the material type was changed, and the primary use was altered from storage to use as a foot pad. Samples were collected over an eight-month period and included three separate flights: Flight 9 (SpX-22; June 28th - July 2nd, 2021), Flight 10 (SpX-23; Sept. 20th - 24th, 2021), and Flight 11 (SpX-24; Jan. 10th - 14th, 2022). During each flight, samples were collected for five consecutive days, using sampling kits that contained sterile gloves and individually packaged sterile pre-wetted polyester wipes: eight for surfaces, one kit control that remained closed, and one environmental control that was removed, unfolded, and waved in the air for 30 seconds. After use, all wipes were transferred to a new sterile bag and stored at 4°C while in orbit and during transport back to the Jet Propulsion Lab (JPL) for processing, no more than five days post-collection. Due to differences in storage temperature requirements or delay in sample return, samples from Mission 1 and Flight 8 of Mission 2 were excluded from community-based metagenomic analyses.

Upon return to Earth, wipes were shaken vigorously for 2 mins in 200 mL of sterile PBS and concentrated with an InnovaPrep CP-150, prior to subdivision into four aliquots: 1500 µL for metagenomics (untreated), 1500 µL for metagenomics (treated), 800 µL for long-term storage, and 200 µL for isolate culturing. Treated metagenomic samples were exposed to propidium monoazide

(PMA), enabling discrimination between viable and dead/compromised cells (i.e., active community and total community potential).

Microbial burden quantification

To quantify the microbial burden, quantitative polymerase chain reaction (qPCR) of the partial 16S rRNA gene (for bacterial quantification) and ITS gene (for fungal quantification) were performed on an Applied Biosystems QuantStudio 6 (Waltham, MA). Zymo's Femto Bacterial DNA Quantification Kit (Cat. # E2006) and Femto Fungal DNA Quantification kit (Cat. # E2007) were used to quantify the 16S rRNA gene and ITS gene respectively. The Femto Bacterial kit included a proprietary degenerate set of primers targeting the 16S rRNA gene, while the Femto Fungal kit used primers ITS-1F and ITS-2R (ITS-1F: 5'- CTTGGTCATTTAGAGGAAGTAA-3', ITS-2R: 5'- GCTGCGTTCTTCATCGATGC-3'). Each reaction consisted of 18 µL of Femto Kit Master Mix (including primers) and 2 µL of template DNA (PMA treated and untreated samples). Samples were run in triplicate, and the average and standard deviation were calculated. Elution fluid from the Maxwell RSC Cell DNA Kit (Promega, Madison, WI) was used as the negative control in each run. The reaction conditions were: a 10-min denaturation at 95 °C, followed by 40 cycles of denaturation at 95 °C for 30 sec, annealing at 50 °C for 30 sec, and extension at 72°C for 1 min. Samples were quantified based on ng of DNA per wipe. The qPCR efficiency was ~ 98% for each run. Negative control values were not deducted, since the values were at ~ 100 copies per 1 or 10 µL and therefore were not scalable. Resulting values (ng/wipe) were log transformed, and differences between sample types were compared with Wilcoxon rank-sum tests, using a Benjamini-Hochberg correction for multiple comparisons.

Metagenomic data accession, sequencing, processing, and analysis

Metagenomic data from Mission 2 is publicly available in the NASA's GeneLab Database, GLDS-252 (https://genelab-data.ndc.nasa.gov/genelab/accession/GLDS-252/), as well as in the National Center for Biotechnology Information (NCBI) Sequence Read Archive (SRA), PRJNA781277 (https://www.ncbi.nlm.nih.gov/search/all/?term=PRJNA781277) (*2*). Newly collected samples were submitted to the Microbiome Core at UC San Diego for DNA extraction and library preparation, following standard Earth Microbiome Project (EMP) protocols (*3*). Samples were then sequenced at the UC San Diego IGM Genomics Center on an Illumina NovaSeq 6000 with positive

(Zymo mock communities) and negative controls (extraction and PCR). To ensure consistency in pre-processing methods, raw data files from Missions 2 and 3 were combined and processed through the same bioinformatic pipeline. Quality was checked with fastqc/multiqc, prior to removal of low quality reads and adapter trimming with fastp (*4–6*). All human-associated reads and control sequences were then mapped to the Telomere-to-Telomere + Y chromosome (T2T-Y) database, using minimap2 (v. 2.28-r1209) (*7–9*). Taxonomy was assigned with metaphlan4 (v. 4.1.1.) (*10*, *11*), and the species-level genome bins (SBG) file was then imported into an R environment (v. 2023.06.0+421) for subsequent community-based analyses and figure generation, using the phyloseq and ggplot2 packages (*12*, *13*). With the exception of the differential abundance analysis (for which the unrarefied data were used), samples were then rarefied to 10,000 reads per sample (loss of $n = 23$ samples) and transformed using a centered log-ratio (CLR) transformation.

Differences in community composition were calculated with Aitchison distance, and compared using permutational multivariate analysis of variance (PERMANOVA) tests. Further, to determine how quickly microbial communities change over time between surfaces, volatility was calculated as the mean within-group Aitchison distance (± SEM) between all samples, with untreated and PMA-treated samples considered separately.

To gain a deeper understanding of fine-scale taxonomic changes, differential abundance (DA) analyses were performed to identify taxa with significant shifts in relative abundance between missions, flights, days of sample collection within flights, and locations. These results were cross-validated using Random Forest models to assess the importance of individual taxa in distinguishing between variables of interest. DA analysis was conducted on unrarefied data using ALDEx2 (*14*), as compared to the Cupola. The Cupola was selected as the reference location, because it represents the most ecologically 'baseline' environment among the sampled locations, with minimal direct or transient microbial inputs. In contrast, other sites such as the WHC, dining table, or crew quarters might be expected to be enriched with microbes from specific external inputs (fecal, food-associated, or human-associated taxa, respectively). To reduce false positives, taxa with an abundance of less than 1e-5% were removed prior to analysis. Models were run with decom = iqlr, and taxa with a p-value $< 0.05$ were retained. DA results were then validated with Random Forest supervised learning models to predict how well each sample was classified using the metadata criteria. Models were run in the randomForest R package, with 70% of samples used for training, 2500 trees per model, and 10-fold cross validation (*15*).

Additionally, a source-tracking analysis was conducted, for which 25 skin-associated taxa (Table S6) were identified and differences in relative abundance between sample types were tested with Wilcoxon rank-sum tests, using a Benjamini-Hochberg correction for multiple comparisons. Any skin-associated taxa that are also commonly found in environmental samples (e.g., *Acinetobacter*) were excluded from this analysis.

Functional annotation and genomic profiling

To quantify traits associated with pathogenic potential, the reads were assembled with MEGAHIT (v. 1.2.9) (*16*), and genes were identified from contigs with ABRicate (Seemann T, Abricate, https://github.com/tseemann/abricate), using the MEGARes database and the VFDB (*17–19*). AMR was characterized to class and mechanism of resistance (e.g., confers resistance to Tetracycline) levels, and data were normalized by millions of sequences per sample, as calculated after filtering, trimming, and host DNA removal. Virulence factors were normalized by contig length (Mbp). Differences between missions were compared with Kruskal-Wallis, and differences between flights, days, and surfaces were tested with Wilcoxon rank-sum tests, using a Benjamini-Hochberg correction for multiple comparisons.

LGT events were estimated with the WAAFLE software package (v. 1.1.0) (*20*), using the standard pipeline. Briefly, genes were mapped to the WAAFLE database and identified as candidate LGT genes. Junction support was then quantified and the data were QC filtered, using a junction score of $\geq$80, support bases $\geq$100, and a confidence classification of $\geq$90%. Resulting files were normalized based on contig length (LGTs/Mbp).

Isolate culturing, sequencing, and functional profiling

For Mission 3, one hundred microliters of aliquot was plated on Reasoner's 2A agar (R2A), potato dextrose agar (PDA), and blood agar to capture environmental bacteria, environmental fungi, and pathogenic bacteria, respectively at three different concentrations ($10^{-1}$, $10^{-2}$, $10^{-3}$). Additionally, 1 mL of aliquot at these same concentrations were plated onto Easy Plate AC (Kikkoman, product #: 61973) and Easy Plate YM-R (Kikkoman, product #: 61977) (also targeting environmental bacteria and fungi respectively). R2A plates, PDA plates, and Easy Plate AC and YM-R were incubated at 25°C for 7 days, and blood agar plates were incubated at 37°C for 2 days. For Missions 2 & 3, 1 mL of aliquot was plated onto Easy Plate AC, Easy Plate YM-R and Easy Plate CC (targeting coliform bacteria; Kikkoman, product #: 61974), and 100 μL of aliquot was plated onto blood agar, at previously specified concentrations ($10^{-1}$, $10^{-2}$, $10^{-3}$). Easy Plate AC and YM-R were incubated at 25°C for 7 days and Easy Plate CC and blood agar plates were incubated at 37°C for 2 days. For all missions, single colonies were re-streaked on fresh plates to ensure pure culture and incubated at the corresponding temperature and time, prior to being archived in 30% glycerol and stored at − 80°C. Mission 1 isolates were whole genome sequenced (WGS) prior to the start of this study (methods published here: (*21*)) and combined with newly generated whole genome sequencing (WGS) data from the Mission 2 and 3 series. Isolates from Missions 2 and 3 were WGS on an Illumina NextSeq 500 or NovaSeq X Plus (paired end, 2 ×150 bp), resulting in ~3T sequences. Quality was assessed with FastQC (v0.11.7), and adapter trimming and filtering was performed with fastp (v0.20.0) (*4*, *5*). Isolate genomes were then assembled with MEGAHIT (v. 1.2.9), completeness and contamination were checked with CheckM (*16*, *22*), and taxonomy was assigned with the GTDB-Tk database (v. 2.4.0) (*23*, *24*). Using the same methods used for metagenomic analysis, assembled genomes were then annotated for AMR and virulence genes with ABRicate (v. 1.0.1), using the MEGARes (https://github.com/tseemann/abricate) and VFDB (*17*, *19*), as well as with the AMRFinderPlus package (v. 3.12.8; database v. 2024-07-22.1). Genes were retained for analysis/interpretation of results if there was > 80% coverage and identity.

For phenotypic profiling, 102 isolates were selected based on taxonomic classification, location of collection, and the abundance of antimicrobial resistance or virulence genes. Taxonomically diverse bacteria were selected with repetition at the species level to allow for both inter- and intra-species comparative analysis (e.g., six strains of *P. fulva* from three locations were tested from Mission 2. Although some strains were identified as good candidates for either AMR or virulence testing independently, there were 44 strains shared between the experiments (Tables 1 and 2).

*In vitro* antibiotic resistance assays

For AMR, 67 strains from 33 representative species were tested, including 24 strains from Mission 1, 21 from Mission 2, and 22 from Mission 3 (Table 1). TSB was inoculated with each isolate and grown overnight in a shaking incubator at 37°C, 180 RPM. 100 μL of the culture was pipetted onto Mueller-Hinton (MH) agar and spread with plating beads. Excess liquid was evaporated prior to experimentation to ensure adherence of the antibiotic discs. Single-disc diffusion was performed following the Kirby-Bauer method (*25*, *26*). Eighteen different antibiotics were tested from four main classes, including four subclasses of β-lactams (Amoxicillin/Clavulanic Acid, Aztreonam, Cefoxitin, Ceftriaxone, Cefazolin, Cloxacillin, Meropenem, Oxacillin, and Penicillin), Fluoroquinolones (Ciprofloxacin, Gemifloxacin, Levofloxacin, Moxifloxacin, Norfloxacin, and Ofloxacin), a Lacosamide (Clindamycin), and Macrolides (Azithromycin and Erythromycin). All antibiotics were tested using premade discs, with the exception of Aztreonam, Penicillin, Cloxacillin, and Gemifloxacin. For these, each antibiotic was inoculated onto blank Oxoid discs, according to the manufacturer's protocols. Plates were incubated at either 37°C or room temperature, based upon the optimal growing temperature of each strain as determined by preliminary testing. After 24 hours, the zone of inhibition (ZOI) was measured and recorded. Each antibiotic was tested in duplicate for all strains, and the ZOI for each replicate was averaged. Susceptibility was determined based on the Clinical & Laboratory Standards Institute (CLSI) guidelines, and identified as belonging to one of three categories: Susceptible, Intermediate, or Resistant, where intermediate levels of resistance indicate potential susceptibility of the antibiotic at higher doses.

False positives and false negatives (i.e., bacteria that were resistant to an antibiotic where no genes were identified that would confer that resistance) were identified. False negatives were tested further via plasmid isolation with the QIAprep Spin Miniprep Kit (QiAgen 2020). Plasmid presence was verified with a Picogreen Assay and Gel Electrophoresis. If no plasmid was detected, the bacterial isolate was considered a true false negative.

*In vivo* virulence testing

To assess virulence, 79 strains from 22 representative species were tested, including 22 strains from Mission 1, 25 from Mission 2, and 32 from Mission 3 (Table 2). Isolates were grown in LB overnight in a shaking incubator at 37°C, 180 RPM. Optical density measurements were taken following the incubation period at a wavelength of 600 nm. Appropriate dilutions were made using PBS to ensure samples read between 0.1 - 0.2 (8x107 - 1.6x108 cells/ml). 35 µl of culture was spread onto 6 x 35 mm petri dishes containing Fasting Killing Assay (FKA) media (10 g NaCl, 17 g BD Bacto™ Dehydrated Agar, 10 g Bactopeptone, 0.1198 g CaCl2, 0.120366 g MgSO4, 3.40215 g KH2PO4, 10 g 20% glucose, 27.32 g D-sorbital/1L DI water) and incubated at 37°C until minimal growth could be observed under an optical microscope. *Pseudomonas aeruginosa* PA14 was used as a positive control (*27*), with *E. coli* OP50 remaining the negative control (*28*). Wildtype N2 Bristol *C. elegan*s (obtained from the *Caenorhabditis* genetics center (CGC), University of Minnesota, USA) were maintained on nematode growth media (NGM), inoculated with *E. coli* OP50 (*29*). Nematode populations were maintained by transferring five (5) L4 worms every three (3) days to avoid overpopulation, nutrient deprivation, and dauer formation (*30*). Prior to experimentation, worm synchronization was performed by transferring 30 adult worms onto sterile NGM plates. Worms were left for 4 - 6 hours to lay eggs before removal. Eggs were left to hatch at 21°C, where synchronized progeny were raised on OP50 for 3 days until worms had reached L3-L4 stage, washed and transferred onto experimental plates containing +/- controls and ISS isolates.

Synchronized worms on sterile NGM plates were washed with 3 ml of M9 buffer to allow for transfer and left on ice for 5 mins so that worms could be extracted from the bottom of the tube. Worms were transferred into new tubes containing M9 and left on ice for 10 mins. This step was repeated once more, before nematodes were washed in Kanamycin (100 ug/ml) solution. Synchronized *C. elegans* were left to bathe in the Kanamycin solution at 21°C for 45 min to remove residual *E.coli* OP50 from the nematode bodies. On completion of the final wash, 10 worms were transferred onto the experimental plates. To keep the worms in the experimental area, 100% glycerol was applied to the plate rim (*31*). *C.elegans* were then counted every hour for 5 hours and at 24, 30 and 48 hours. At each timepoint, dead worms were removed and the ratio of confirmed alive:confirmed dead was recorded (two replicates for each space-flown isolate and +/- controls) (*29–31*).

Differences in killing capacity, compared to positive and negative controls, was then visualized and quantified with the online application for survival analysis 2 (OASIS 2) interface, where differences in mean lifespan were compared with Log-Rank Tests (LRTs), using a Bonferroni correction for multiple comparisons, and differences in survival over time were compared with Fisher's Exact Tests (FETs) (*32*). FETs were calculated at four time intervals, over the duration of the testing period (at 25%, 50%, 75%, and 90%). Based on the LRTs and FETs at 90%, bacteria were grouped into one of three categories, indicating virulence potential; avirulent, intermediate virulence, and highly virulent. Avirulent is defined as bacterial strains that had no observable or minimal rates of killing (no statistical difference from *E. coli*). Intermediate virulence were those bacterial strains that showed moderate rates of killing (significantly higher than *E. coli* but significantly less than *P. aeruginosa*), and bacteria were considered highly virulent if the rate of killing was equal to or greater than that of *P. aeruginosa* (no statistical difference from *P. aeruginosa* or significantly higher than *P. aeruginosa*). For bacteria characterized as highly virulent, FETs were assessed at all time intervals to determine the approximate time at which the rate of killing was greater than the positive control and if this trend was consistent over the entire 24-hour monitoring period. Further, highly virulent strains were tested (in duplicate, as above) on three separate occasions to confirm results and then assessed for classification as a false positive or false negative negative result. Strain was considered a false positive if there were greater than 20 VFs identified with an avirulent or intermediate phenotype and false negative if there were no VFs identified with intermediate or highly virulent phenotype.

Metagenome-assembled genome (MAG) analysis

The quality of raw reads from each sample was checked with fastqc (v. 0.11.9) and summarized with multiqc v1.14 (*4*, *33*). Further quality filtering criteria were based on these raw read stats using fastp v.0.23.2 (*5*, *6*), with following filtering criteria: --average_qual 30 --length_required 150 --detect_adapter_for_pe --correction --overrepresentation_analysis --low_complexity_filter. Human reads were removed by mapping against T2T-chm13v2 with bowtie (v. 2.4.4) (*7*, *34*) and extracting unmapped reads with samtools v1.17 (*10*, *35*). Reads were assembled into contigs and scaffolds with MEGAHIT (v. 1.2.3) and its meta-sensitive preset (*16*, *36*). Metagenome-atlas (v. 2.18.1) (*37*) snakemake pipeline was used for genome binning with Metabat2 (*38*), MaxBin2 (*39*), and scoring bins with DASTool (*40*) in the end (as specified in the respective config.yaml file of the pipeline). Resulting bins were further dereplicated with dRep (v. 3.0.0) and fastANI (*41*, *42*). Representative genomes (MAGs, metagenome assembled genomes) were classified with GTDBtk (v. 2.4.0), using reference data version r220 (*24*, *43*). Functional and metabolic annotation was performed with DRAM (v. 1.5.0) and eggNOG mapper (v. 2.1.8) (*44*, *45*). Contigs and bins were evaluated and profiled for their gene content with MetaQUAST (v. 5.2.0) (*46*). Quantification of MAGs was achieved at a minimum coverage of 80% with coverM v0.6.1 (*47*), and growth rate estimations were conducted with GRiD v1.3 in multiplex mode with a minimum coverage cutoff of 0.2, allowing reassignment of ambiguous reads with Pathoscope2 (*48*, *49*). Growth conditions of MAGs were determined with GenomeSPOT (v. 1.0.1) (*50*). ABRicate (v. 1.0.1) was used to profile MAGs and contigs for AMR and VFs according to the following databases: ARG-ANNOT,

CARD, EcOH, Ecoli_VF, MEGARES, NCBI AMRFinderPlus, PlasmidFinder, Resfinder and VFDB (*17*, *18*, *51–57*). Strain-level genomics and popANI comparisons were carried out with inStrain (v. 1.9.0) (*58*).

To correct for assembly-related artifacts and to provide a comprehensive view of all domains of life, species profiling on read and contig levels was conducted with Kraken2/Bracken and the PlusPFP database, using the QIIME2 (qiime2-metagenome_2024.10) framework (*59–61*). Furthermore, the QIIME2 analysis covered removal of potential contaminants with decontam, core metrics of alpha and beta diversity, differential abundance with ANCOM-BC, and feature (taxa and functions) volatility with the q2-longitudinal plugin (*62–64*).

For an in-depth comparative genomics analysis 10 genomes were chosen and processed with the MaGe LABGeM MicroScope platform v3.17.5 with default settings (*65*). This comprehensive analysis covered standard genomics (genome statistics: CheckM; minimal gene sets; annotations: GTDBtk, eggNOG; genome map visualizations: CGView), and a plethora of comparative genomic tools (Mash genome clustering and neighbor-joining; gene phyloprofiles according to the PkGDB database; regions of genomic plasticity: RGP Finder, PkGDB database, AlienHunter, SIGI-HMM; conserved synteny: LinePlot; gene fusion and fission according to the PkGDB database; PkGDB synteny and RefSeq synteny statistics with bi-directional best hit criteria; pan -and core genome analysis based on MICFAM gene families using the SiLiX tool; prediction of antibiotic resistances using RGI v5.0.0 and the CARD database v3.0.2; predictions of virulence according to VFDB and VirulenceFinder; predictions of integrons with IntegronFinder v2.0.2; prediction of macromolecular systems using MacSyFinder v2.1.2; prediction of prophages with Phigaro v2.4.0; prediction of defense systems using DefenseFinder v1.2.2 and CRISPRCasFinder v4.3.2; and metabolic predictions including KEGG, MicroCyc, BioCyc, MetaCyc databases, and prediction of secondary metabolite biosynthesis with AntiSMASH v7.1.0.1.

61. E. Bolyen, J. R. Rideout, M. R. Dillon, N. A. Bokulich, C. C. Abnet, G. A. Al-Ghalith, H. Alexander, E. J. Alm, M. Arumugam, F. Asnicar, Y. Bai, J. E. Bisanz, K. Bittinger, A. Brejnrod, C. J. Brislawn, C. T. Brown, B. J. Callahan, A. M. Caraballo-Rodríguez, J. Chase, E. K. Cope, R. Da Silva, C. Diener, P. C. Dorrestein, G. M. Douglas, D. M. Durall, C. Duvallet, C. F. Edwardson, M. Ernst, M. Estaki, J. Fouquier, J. M. Gauglitz, S. M. Gibbons, D. L. Gibson, A. Gonzalez, K. Gorlick, J. Guo, B. Hillmann, S. Holmes, H. Holste, C. Huttenhower, G. A. Huttley, S. Janssen, A. K. Jarmusch, L. Jiang, B. D. Kaehler, K. B. Kang, C. R. Keefe, P. Keim, S. T. Kelley, D. Knights, I. Koester, T. Kosciolek, J. Kreps, M. G. I. Langille, J. Lee, R. Ley, Y.-X. Liu, E. Loftfield, C. Lozupone, M. Maher, C. Marotz, B. D. Martin, D. McDonald, L. J. McIver, A. V. Melnik, J. L. Metcalf, S. C. Morgan, J. T. Morton, A. T. Naimey, J. A. Navas-Molina, L. F. Nothias, S. B. Orchanian, T. Pearson, S. L. Peoples, D. Petras, M. L. Preuss, E. Pruesse, L. B. Rasmussen, A. Rivers, M. S. Robeson 2nd, P. Rosenthal, N. Segata, M. Shaffer, A. Shiffer, R. Sinha, S. J. Song, J. R. Spear, A. D. Swafford, L. R. Thompson, P. J. Torres, P. Trinh, A. Tripathi, P. J. Turnbaugh, S. Ul-Hasan, J. J. J. van der Hooft, F. Vargas, Y. Vázquez-Baeza, E. Vogtmann, M. von Hippel, W. Walters, Y. Wan, M. Wang, J. Warren, K. C. Weber, C. H. D. Williamson, A. D. Willis, Z. Z. Xu, J. R. Zaneveld, Y. Zhang, Q. Zhu, R. Knight, J. G. Caporaso, Reproducible, interactive, scalable and extensible microbiome data science using QIIME 2. *Nat. Biotechnol.* **37**, 852–857 (2019).

62. N. M. Davis, D. M. Proctor, S. P. Holmes, D. A. Relman, B. J. Callahan, Simple statistical identification and removal of contaminant sequences in marker-gene and metagenomics data. *Microbiome* **6**, 226 (2018).

63. H. Lin, S. D. Peddada, Analysis of compositions of microbiomes with bias correction. *Nat. Commun.* **11**, 3514 (2020).

64. N. A. Bokulich, M. R. Dillon, Y. Zhang, J. R. Rideout, E. Bolyen, H. Li, P. S. Albert, J. G. Caporaso, Q2-longitudinal: Longitudinal and paired-sample analyses of microbiome data. *mSystems* **3** (2018).

65. D. Vallenet, A. Calteau, M. Dubois, P. Amours, A. Bazin, M. Beuvin, L. Burlot, X. Bussell, S. Fouteau, G. Gautreau, A. Lajus, J. Langlois, R. Planel, D. Roche, J. Rollin, Z. Rouy, V. Sabatet, C. Médigue, MicroScope: an integrated platform for the annotation and exploration of microbial gene functions through genomic, pangenomic and metabolic comparative analysis. *Nucleic Acids Res.* **48**, D579–D589 (2020).

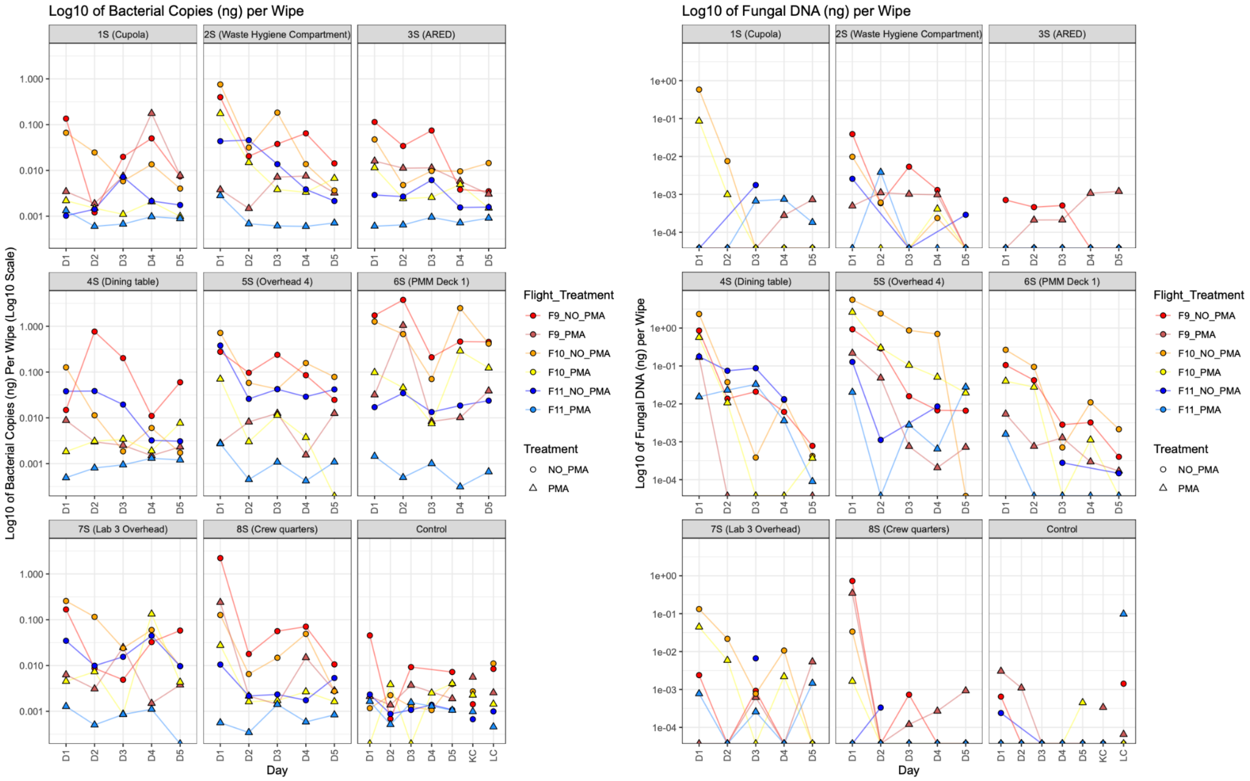

**Fig. S1. Bacterial and fungal qPCR results from Mission 3.** qPCR results include data collected from three flights (F9, F10, and F 11) for both untreated and PMA-treated samples across eight surfaces. Overhead 4 indicates the Stowage rack location. *p-value < 0.05 **p-value < 0.01 ***p-value < 0.001

A

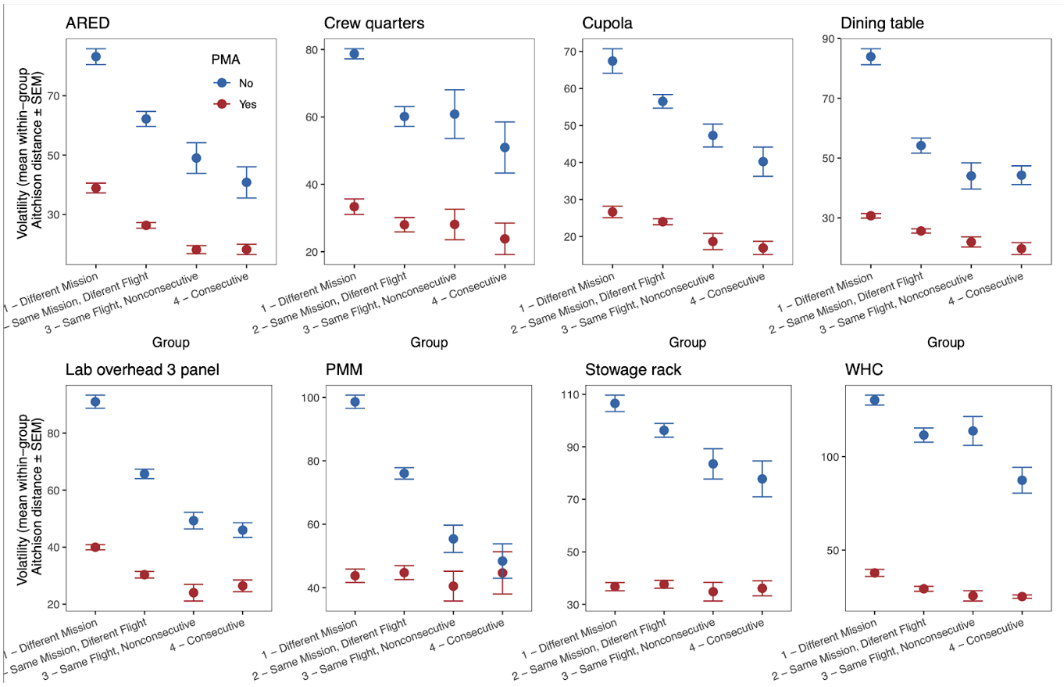

B

| | 1Yv1N | 2Yv2N | 3Yv3N | 4Yv4N | 1Yv2Y | 2Yv3Y | 3Yv4Y | 1Nv2N | 2Nv3N | 3Nv4N |
|---|---|---|---|---|---|---|---|---|---|---|
| ARED | 4.87E-17 | 1.77E-20 | 5.66E-06 | 0.00052024 | 1.92E-09 | 0.00022712 | 0.81252427 | 9.53E-07 | 0.03396774 | 0.31650259 |
| Crew quarters | 1.54E-17 | 3.45E-12 | 0.00139312 | 0.00728344 | 0.02821469 | 0.90694796 | 0.61602921 | 4.51E-06 | 0.81111515 | 0.3240847 |
| Cupola | 4.24E-11 | 2.81E-19 | 5.00E-05 | 0.00044867 | 0.15272954 | 0.06013513 | 0.47572218 | 0.0221188 | 0.06421458 | 0.20059073 |
| Dining table | 1.58E-16 | 5.49E-15 | 0.00082327 | 0.00146006 | 1.70E-05 | 0.06681862 | 0.59146115 | 1.48E-09 | 0.12716832 | 0.96108266 |
| Lab overhead 3 panel | 1.49E-16 | 2.66E-22 | 5.41E-05 | 0.00108744 | 1.72E-07 | 0.02895969 | 0.54954388 | 8.80E-12 | 0.00011434 | 0.13229889 |
| PMM | 5.65E-17 | 9.39E-14 | 0.02116699 | 0.22904867 | 0.94139654 | 0.2361068 | 0.74450526 | 1.71E-10 | 0.00017962 | 0.19657242 |
| Stowage rack | 1.95E-18 | 6.00E-21 | 2.77E-05 | 0.00032828 | 0.98044987 | 0.45270494 | 0.97111464 | 0.02941835 | 0.03833764 | 0.25353781 |
| WHC | 4.16E-17 | 2.98E-18 | 2.08E-05 | 0.00241465 | 0.00026976 | 0.29762147 | 0.95992166 | 0.0004971 | 0.81427619 | 0.04202058 |

**Fig. S2. Volatility of bacterial communities on ISS surfaces, based on PMA treatment at different time scales.** (**A**) Mission represents samples that were collected up to 5 years apart. Different flights represent periods that span months, and consecutive days are samples that were collected one day apart. Volatility was calculated as the mean within-group Aitchison distance (± SEM) between all samples, with untreated and PMA-treated samples considered separately. (**B**) Statistical comparisons between and within treatments over time, as calculated with two-sided Wilcoxon tests. N indicates untreated samples. Y indicates PMA treatment. Number (i.e., 1, 2, 3, and 4) corresponds to the time period represented in panel A.

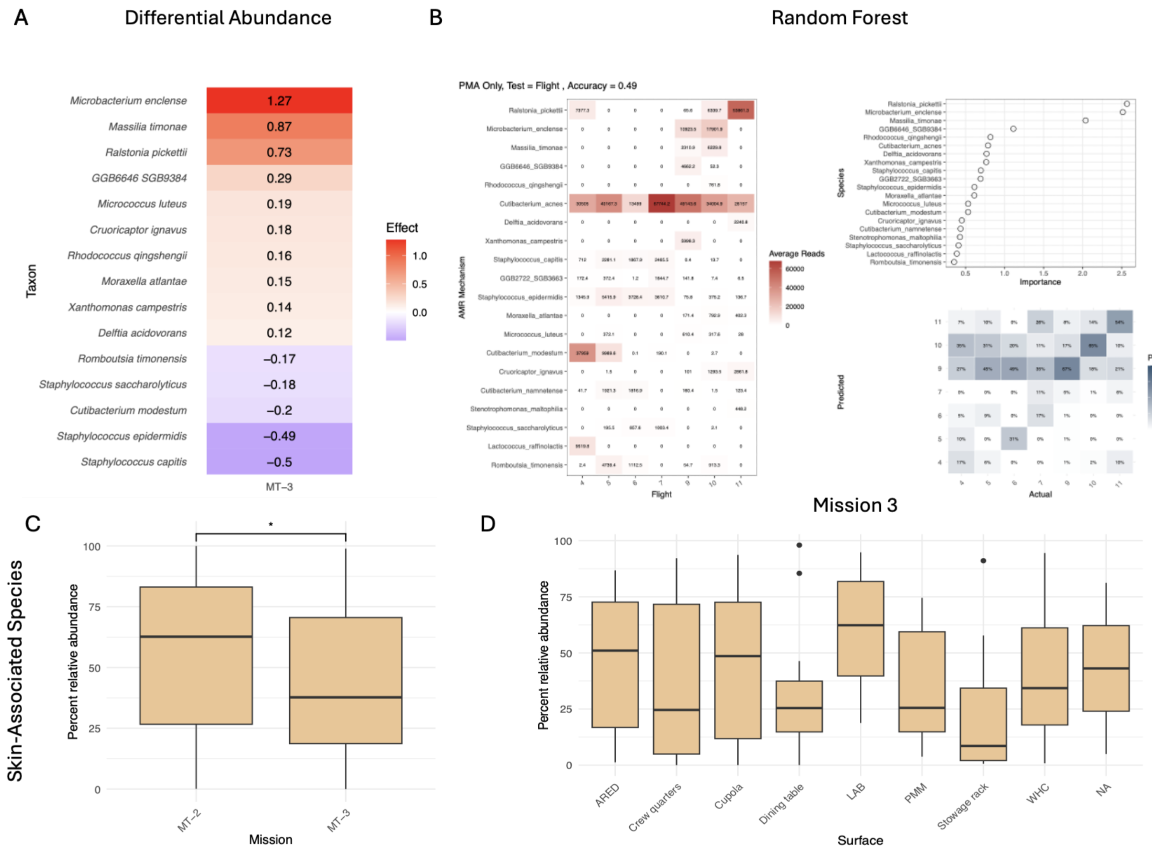

**Fig. S3. Enrichment of different bacterial taxonomic groups over time from viable (PMA-treated) communities collected from the ISS, based on magnitude of change and predictive modeling across all taxa, as well as quantification of skin-associated taxa only.** (**A**) Differential abundance (DA) analysis, representing the log-fold change (LFC) of taxa between Mission 2 and Mission 3 for all surfaces, using ALDEx2. (**B**) We identified the top 20 taxa ranked by feature importance in a Random Forest model that represent the most informative features for distinguishing between flights (accuracy = 0.49, error ratio = 3.4) (**C**) Percent relative abundance of skin-associated taxa identified between missions and among surfaces, based on 25 common skin associates. Though there was variation among samples and a slight decrease in skin-associated taxa over time, there was a general enrichment throughout the ISS, with no statistical differences observed between surfaces (Wilcoxon rank-sum tests, using a Benjamini-Hochberg correction for multiple comparisons).

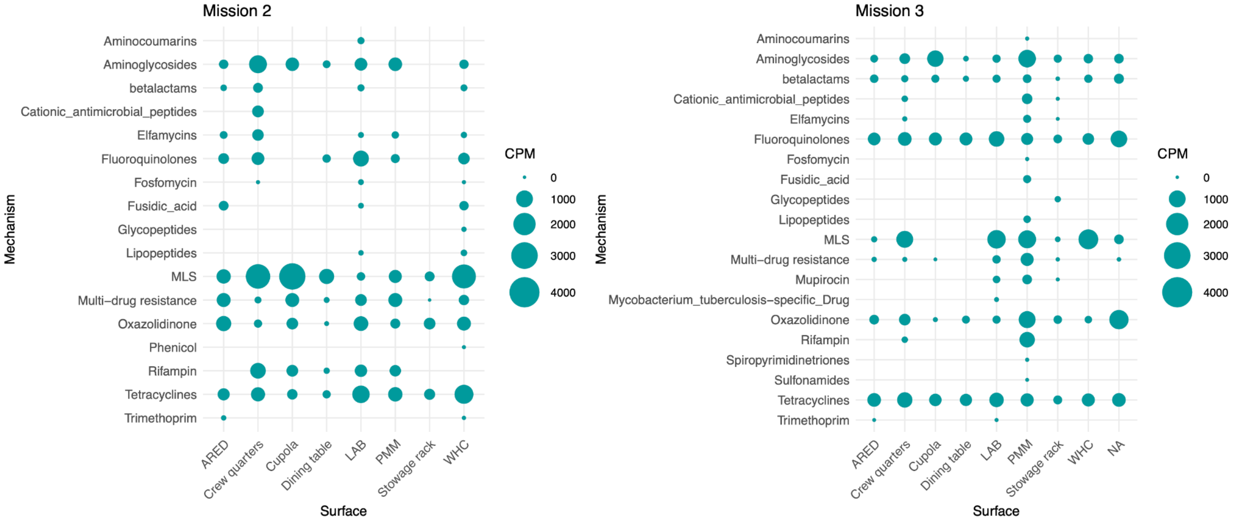

**Fig. S4. Proportion of antimicrobial resistance (AMR) genes recovered from viable communities from surfaces, for each mission.** There were no significant differences observed in the proportion of mechanism-encoded AMR genes between missions or among surfaces.

| **16S, Untreated** | | | | | | | | |
|---|---|---|---|---|---|---|---|---|
| Location | 1S (Cupola) | 2S (WHC) | 3S (ARED) | 4S (Dining table) | 5S (Stowage rack) | 6S (PMM Deck 1) | 7S (Lab 3 Overhead) | 8S (Crew quarters) |
| 2S (WHC) | 0.067 | NA | NA | NA | NA | NA | NA | NA |
| 3S (ARED) | 0.829 | 0.083 | NA | NA | NA | NA | NA | NA |
| 4S (Dining table) | 0.300 | 0.464 | 0.300 | NA | NA | NA | NA | NA |
| 5S (Stowage rack) | < 0.001*** | 0.079 | 0.001** | 0.017* | NA | NA | NA | NA |
| 6S (PMM Deck 1) | < 0.001*** | 0.028* | < 0.001*** | 0.007** | 0.389 | NA | NA | NA |
| 7S (Lab 3 Overhead) | 0.033* | 1.000 | 0.028* | 0.503 | 0.045* | 0.016* | NA | NA |
| 8S (Crew quarters) | 0.389 | 0.389 | 0.426 | 0.755 | 0.009** | 0.005** | 0.310 | NA |
| Control | 0.028* | < 0.001*** | 0.006** | 0.001** | < 0.001*** | < 0.001*** | < 0.001*** | < 0.001*** |

| **16S, PMA Treated** | | | | | | | | |
|---|---|---|---|---|---|---|---|---|
| Location | 1S (Cupola) | 2S (WHC) | 3S (ARED) | 4S (Dining table) | 5S (Stowage rack) | 6S (PMM Deck 1) | 7S (Lab 3 Overhead) | 8S (Crew quarters) |
| 2S (WHC) | 0.875 | NA | NA | NA | NA | NA | NA | NA |
| 3S (ARED) | 0.875 | 0.962 | NA | NA | NA | NA | NA | NA |
| 4S (Dining table) | 0.930 | 0.783 | 0.875 | NA | NA | NA | NA | NA |

| | | | | | | | | |
|---|---|---|---|---|---|---|---|---|
| 5S (Stowage rack) | 0.875 | 0.930 | 0.962 | 0.875 | NA | NA | NA | NA |
| 6S (PMM Deck 1) | 0.535 | 0.535 | 0.535 | 0.535 | 0.535 | NA | NA | NA |
| 7S (Lab 3 Overhead) | 0.930 | 0.930 | 0.962 | 0.930 | 1.000 | 0.535 | NA | NA |
| 8S (Crew quarters) | 0.962 | 0.783 | 0.925 | 0.930 | 0.930 | 0.535 | 0.930 | NA |
| Control | 0.962 | 0.535 | 0.535 | 0.875 | 0.579 | 0.535 | 0.783 | 0.930 |
| **ITS, Untreated** | | | | | | | | |
| Location | 1S (Cupola) | 2S (WHC) | 3S (ARED) | 4S (Dining table) | 5S (Stowage rack) | 6S (PMM Deck 1) | 7S (Lab 3 Overhead) | 8S (Crew quarters) |
| 2S (WHC) | 0.188 | NA | NA | NA | NA | NA | NA | NA |
| 3S (ARED) | 0.730 | 0.027* | NA | NA | NA | NA | NA | NA |
| 4S (Dining table) | 0.004** | 0.005** | < 0.001*** | NA | NA | NA | NA | NA |
| 5S (Stowage rack) | 0.004** | 0.005** | < 0.001*** | 0.420 | NA | NA | NA | NA |
| 6S (PMM Deck 1) | 0.011* | 0.067 | 0.001** | 0.282 | 0.067* | NA | NA | NA |
| 7S (Lab 3 Overhead) | 0.189 | 0.649 | 0.023* | 0.041* | 0.022* | 0.294 | NA | NA |
| 8S (Crew quarters) | 0.768 | 0.269 | 0.547 | 0.005** | 0.005** | 0.017* | 0.250 | NA |
| Control | 0.583 | 0.016* | 0.828 | < 0.001*** | < 0.001*** | < 0.001*** | 0.016* | 0.390 |
| **ITS, PMA** | | | | | | | | |

| Treated | | | | | | | | |
|---|---|---|---|---|---|---|---|---|
| Location | 1S (Cupola) | 2S (WHC) | 3S (ARED) | 4S (Dining table) | 5S (Stowage rack) | 6S (PMM Deck 1) | 7S (Lab 3 Overhead) | 8S (Crew quarters) |
| 2S (WHC) | 0.963 | NA | NA | NA | NA | NA | NA | NA |
| 3S (ARED) | 0.476 | 0.536 | NA | NA | NA | NA | NA | NA |
| 4S (Dining table) | 0.388 | 0.265 | 0.111 | NA | NA | NA | NA | NA |
| 5S (Stowage rack) | 0.007** | 0.006** | 0.001** | 0.187 | NA | NA | NA | NA |
| 6S (PMM Deck 1) | 0.438 | 0.367 | 0.126 | 0.658 | 0.044* | NA | NA | NA |
| 7S (Lab 3 Overhead) | 0.601 | 0.487 | 0.187 | 0.536 | 0.025* | 0.887 | NA | NA |
| 8S (Crew quarters) | 0.658 | 0.749 | 0.749 | 0.224 | 0.006** | 0.291 | 0.420 | NA |
| Control | 0.536 | 0.658 | 0.822 | 0.126 | 0.001** | 0.187 | 0.287 | 0.887 |

**Table S1. Statistical results for qPCR results.** Bacterial (16S rRNA gene) and fungal (ITS region), as well as untreated and PMA-treated, samples were considered separately. Differences between surfaces were calculated with Wilcoxon rank-sum tests, using a Benjamini-Hochberg correction for multiple comparisons. *p-value < 0.05 **p-value < 0.01 ***p-value < 0.001

| Comparison 1. Differences between flights. | | | | | | | | |
|---|---|---|---|---|---|---|---|---|
| | | | ARG | | VF | | LGT | |
| Mission | FlightA | FlightB | p-value | Significance | p-value | Significance | p-value | Significance |
| Mission 2 | 4 | 5 | 0.312 | ns | 0.67 | ns | 0.962 | ns |
| Mission 2 | 4 | 6 | 0.016 | * | 0.67 | ns | 0.962 | ns |
| Mission 2 | 4 | 7 | 0.197 | ns | 0.67 | ns | 0.962 | ns |
| Mission 2 | 5 | 6 | 0.003 | ** | 0.31 | ns | 0.962 | ns |
| Mission 2 | 5 | 7 | 0.047 | * | NA | NA | 0.962 | ns |
| Mission 2 | 6 | 7 | 0.916 | ns | 0.31 | ns | 0.965 | ns |
| Mission 3 | 9 | 10 | 0.916 | ns | 0.98 | ns | 0.962 | ns |
| Mission 3 | 9 | 11 | 0.01 | ** | 0.67 | ns | 0.730 | ns |
| Mission 3 | 10 | 11 | 0.015 | * | 0.67 | ns | 0.506 | ns |
| Comparison 2. Differences between consecutive sampling days. | | | | | | | | |
| | | | ARG | | VF | | LGT | |
| Mission | DayA | DayB | p-value | Significance | p-value | Significance | p-value | Significance |
| Mission 3 | 1 | 2 | 0.119 | ns | 0.429 | ns | 0.322 | ns |
| Mission 3 | 1 | 3 | 0.575 | ns | NA | NA | 0.749 | ns |
| Mission 3 | 1 | 4 | 0.367 | ns | 0.558 | ns | 0.990 | ns |
| Mission 3 | 1 | 5 | 0.879 | ns | NA | NA | 0.520 | ns |
| Mission 3 | 2 | 3 | 0.367 | ns | NA | NA | 0.520 | ns |
| Mission 3 | 2 | 4 | 0.037 | * | 0.558 | ns | 0.433 | ns |
| Mission 3 | 2 | 5 | 0.119 | ns | NA | NA | 0.749 | ns |
| Mission 3 | 3 | 4 | 0.119 | ns | NA | NA | 0.822 | ns |
| Mission 3 | 3 | 5 | 0.672 | ns | NA | NA | 0.749 | ns |
| Mission 3 | 4 | 5 | 0.218 | ns | NA | NA | 0.749 | ns |
| Comparison 3. Differences between surface type. | | | | | | | | |
| | | | ARG | | VF | | LGT | |

| Mission | Surface A | SurfaceB | p-value | Significance | p-value | Significance | p-value | Significance |
|---|---|---|---|---|---|---|---|---|
| Mission 3 | ARED | Crew quarters | NA | NA | NA | NA | 0.021 | * |
| Mission 3 | ARED | Cupola | NA | NA | NA | NA | 0.911 | ns |
| Mission 3 | ARED | Dining table | NA | NA | NA | NA | 0.114 | ns |
| Mission 3 | ARED | Lab 3 | NA | NA | NA | NA | 0.802 | ns |
| Mission 3 | ARED | PMM | NA | NA | NA | NA | 0.403 | ns |
| Mission 3 | ARED | Stowage rack | NA | NA | NA | NA | 0.851 | ns |
| Mission 3 | ARED | WHC | NA | NA | NA | NA | 0.462 | ns |
| Mission 3 | Crew quarters | Cupola | NA | NA | NA | NA | 0.021 | * |
| Mission 3 | Crew quarters | Dining table | 0.758 | ns | 1 | ns | 0.212 | ns |
| Mission 3 | Crew quarters | Lab 3 | 0.157 | ns | 0.709 | ns | 0.013 | * |
| Mission 3 | Crew quarters | PMM | 0.005 | ** | 0.709 | ns | 0.005 | ** |
| Mission 3 | Crew quarters | Stowage rack | NA | NA | NA | NA | 0.005 | ** |
| Mission 3 | Crew quarters | WHC | NA | NA | NA | NA | 0.005 | ** |
| Mission 3 | Cupola | Dining table | NA | NA | NA | NA | 0.147 | ns |
| Mission 3 | Cupola | Lab 3 | NA | NA | NA | NA | 0.532 | ns |
| Mission 3 | Cupola | PMM | NA | NA | NA | NA | 0.212 | ns |
| Mission 3 | Cupola | Stowage rack | NA | NA | NA | NA | 0.942 | ns |
| Mission 3 | Cupola | WHC | NA | NA | NA | NA | 0.212 | ns |
| Mission 3 | Dining table | Lab 3 | 0.028 | * | 0.709 | ns | 0.109 | ns |
| Mission 3 | Dining table | PMM | 0.003 | ** | 0.709 | ns | 0.021 | * |
| Mission 3 | Dining table | Stowage rack | NA | NA | NA | NA | 0.029 | * |

| | | | | | | | | |
|---|---|---|---|---|---|---|---|---|
| Mission 3 | Dining table | WHC | NA | NA | NA | NA | 0.016 | * |
| Mission 3 | Lab 3 | PMM | 0.028 | * | 0.709 | ns | 0.532 | ns |
| Mission 3 | Lab 3 | Stowage rack | NA | NA | NA | NA | 0.501 | ns |
| Mission 3 | Lab 3 | WHC | NA | NA | NA | NA | 0.445 | ns |
| Mission 3 | PMM | Stowage rack | NA | NA | NA | NA | 0.109 | ns |
| Mission 3 | PMM | WHC | NA | NA | NA | NA | 0.802 | ns |
| Mission 3 | Stowage rack | WHC | NA | NA | NA | NA | 0.212 | ns |
| Comparison 4. Differences among individual surface types over time. | | | | | | | | |
| | | | **ARG** | | **VF** | | **LGT** | |
| **Surface** | **Mission A** | **Mission B** | **p-value** | **Significance** | **p-value** | **Significance** | **p-value** | **Significance** |
| ARED | Mission 2 | Mission 3 | 0.366 | ns | 0.082 | ns | 0.205 | ns |
| Crew quarters | Mission 2 | Mission 3 | 0.417 | ns | 0.437 | ns | 0.506 | ns |
| Cupola | Mission 2 | Mission 3 | 0.954 | ns | NA | NA | 0.182 | ns |
| Dining table | Mission 2 | Mission 3 | 0.217 | ns | 0.331 | ns | 0.541 | ns |
| Lab 3 | Mission 2 | Mission 3 | 1 | ns | 0.567 | ns | 0.956 | ns |
| PMM | Mission 2 | Mission 3 | 0.044 | * | NA | NA | 1 | ns |
| Stowage rack | Mission 2 | Mission 3 | 0.221 | ns | 0.349 | ns | 0.788 | ns |
| WHC | Mission 2 | Mission 3 | 0.461 | ns | NA | NA | 0.064 | ns |

**Table S2. Statistical comparisons for differences in antimicrobial resistance gene (AMR), virulence factor (VF) gene, and lateral gene transfer (LGT) event proportion from metagenomic data for PMA-treated samples.** Comparisons include (1) differences between flights, (2) consecutive sampling days, (3) surface type (e.g., ARED compared to the Cupola), and (4) among individual surfaces types over time (e.g., ARED between missions). Flights were only compared within missions, and consecutive sampling days are considered within each flight (e.g., day 1 compared to day 2 within flight 9). Comparisons (2) and (3) are for MT-3 only, as MT-3 is the only mission that had consecutive sampling days, and differences between surfaces were already characterized from the Mission 2 study. AMR abundance was normalized by millions of sequences per sample (CPM), VFs were normalized by contig length (VFs/Mbp), and LGT events were normalized by LGT events/Mbp. Untreated and PMA-treated samples were considered separately. P-values calculated with Wilcoxon rank-sum tests, using a Benjamini-Hochberg correction for multiple comparisons. 'NA' indicates zero values. *p-value < 0.05 **p-value < 0.01 ***p-value < 0.001

[Uploaded as attachment, due to size]

**Table S3. Isolate antimicrobial resistance (AMR) assay results.** Performed with Kirby-Bauer tests, in which each isolate (n = 67) was tested for resistance to 18 antibiotics from four main classes, including nine antibiotics from four subclasses of β-lactams, six Fluoroquinolones, one Lacosamide, and two Macrolides. False positive results (for which an encoding gene was present, but there was no resistance observed) are highlighted in blue. False negative results (for which there was no gene identified, but for which there was resistance observed) are highlighted in green.

[Uploaded as attachment, due to size]

**Table S4. Isolate virulence testing results.** Virulence was tested in a *C. elegans* model (n = 79 isolates), where percent survival was averaged across replicates, prior to calculation of differences in mean lifespan and survival over time, compared to *E. coli* OP50 (EC; negative control) and *P. aeruginosa* PA14 (PA; positive control). Results were considered a false positive if there were greater than 20 virulence factors (VFs) identified with an avirulent or intermediate phenotype and a false negative if there were no VFs identified with an intermediate or highly virulent phenotype. *p-value < 0.05 **p-value < 0.01 ***p-value < 0.001

| Abbreviation | Location description | ISS module |
|---|---|---|
| ARED | Advanced resistive exercise device foot platform | Node 3 |
| Crew quarters | Port crew quarters, bump out exterior aft wall | Node 2 |
| Cupola | Port panel next to the cupola | Node 3 |
| Dining table | Dining table | Node 1 |
| LAB | Panel of the Materials Science Research Rack 1 (MSRR-1) | LAB |
| PMM | Permanent multipurpose module | Port 1 |
| Stowage rack | Overhead 4 (Zero G stowage rack) | Node 1 |
| WHC | Waste and hygiene compartment | Node 3 |

**Table S5. Description of sample locations sampled on the ISS.** Abbreviation corresponds to abbreviation used throughout the manuscript. For Mission 3 (MT-3), each location was sampled for five consecutive days for three flights (9, 10, and 11), from June 2021 to January 2022.

| **Taxon** |
|---|
| *Corynebacterium accolens* |
| *Corynebacterium amycolatum* |
| *Corynebacterium jeikeium* |
| *Corynebacterium kroppenstedtii* |
| *Corynebacterium mucifaciens* |
| *Corynebacterium striatum* |
| *Corynebacterium tuberculostearicum* |
| *Cutibacterium acnes* |
| *Cutibacterium avidum* |
| *Cutibacterium granulosum* |
| *Cutibacterium modestum* |
| *Cutibacterium namnetense* |
| *Finegoldia magna* |
| *Lawsonella clevelandensis* |
| *Micrococcus luteus* |
| *Peptoniphilus spp.* |
| *Staphylococcus aureus* |
| *Staphylococcus capitis* |
| *Staphylococcus epidermidis* |
| *Staphylococcus haemolyticus* |
| *Staphylococcus hominis* |
| *Staphylococcus lugdunensis* |
| *Staphylococcus simulans* |
| *Staphylococcus warneri* |
| *Staphylococcus xylosus* |

**Table S6. Skin-associated taxa.** Taxa selected only if primarily found to be associated with human skin.
